# 4.1N and SAP97 regulate different phases of AMPA receptor intracellular transport

**DOI:** 10.1101/2022.09.05.506328

**Authors:** Caroline Bonnet, Justine Charpentier, Natacha Retailleau, Daniel Choquet, Françoise Coussen

## Abstract

Changes in the number of synaptic AMPA subtypes of glutamate receptors (AMPAR) underlie many forms of synaptic plasticity. These variations are controlled by a complex interplay between their intracellular transport (IT), export to the plasma membrane, stabilization at synaptic sites, and recycling. The differential molecular mechanisms involved in these various trafficking pathways and their regulation remains partly unknown. We have recently reported the visualization of AMPAR IT in cultured hippocampal neurons and demonstrated its regulation during synaptic plasticity inducing protocols (Hangen, Cordelieres et al., 2018), opening the path to the differential analysis of the mechanisms controlling AMPAR transport and exocytosis.

The cytosolic C-terminal (C-ter.) domain of AMPAR GluA1 subunit is specifically associated with cytoplasmic proteins that could be implicated in the regulation of their IT such as 4.1N and SAP97. Here we analyze how interactions between GluA1 and 4.1N or SAP97 regulate IT and exocytosis at the plasma membrane in basal condition and after cLTP induction. We use sh-RNA against 4.1N and SAP97 and specific mutations and deletions of GluA1 C-ter. domain to characterize how these interactions are involved in coupling AMPAR to the transport machinery.

The down-regulation of both 4.1N or SAP97 by shRNAs decrease GluA1 containing vesicle number, modify their transport properties and decrease GluA1 export to the PM, indicating their role in GluA1 IT. The total deletion of the C-ter. domain of GluA1 fully suppresses its IT. Disruption of GluA1 binding to 4.1N decreases the number of GluA1 containing transport vesicles, inhibits GluA1 externalization but does not affect the transport properties of the remaining GluA1 containing vesicles. This indicates a role of the 4.1N-GluA1 interaction during exocytosis of the receptor in basal transmission. In contrast, disrupting the binding between SAP97 and GluA1 modifies the basal transport properties of GluA1 containing vesicles and decreases GluA1 export to the plasma membrane. Importantly, disrupting GluA1 interaction with either 4.1N or SAP97 prevents both the cLTP induced increase in the number of GluA1 containing vesicles observed in control and GluA1 externalization. Our results demonstrate that specific interactions between 4.1N or SAP97 with GluA1 have different roles in GluA1 IT and exocytosis. During basal transmission, the binding of 4.1N to GluA1 allows the fusion/fission membrane exocytosis whereas the interaction with SAP97 is essential for GluA1 IT. During cLTP the interaction of 4.1N with GluA1 allows both IT and exocytosis of the receptor in hippocampal cultured neurons. Altogether, our results identify the differential roles of 4.1N and SAP97 in the control of various phases of GluA1 IT.

## Introduction

AMPA-type glutamate receptors (AMPAR) are ionotropic tetrameric receptors activated by glutamate, the main excitatory neurotransmitter of the central nervous system. The synaptic targeting, clustering, and immobilization of glutamate receptors in the post-synaptic density in front of glutamate release sites are crucial for efficient excitatory synaptic transmission. The number of neurotransmitter receptors, and particularly AMPAR, present at the synapse is regulated by a complex set of interdependent mechanisms going from biogenesis, intracellular transport (IT) (Diaz-Alonso & Nicoll,2021, Hiester, Becker et al., 2018), externalization at the plasma membrane (PM), lateral diffusion (Choquet & Triller, 2013), stabilization at synapses and trafficking in and out synaptic sites (Groc & Choquet, 2020). In highly polarized and spatially extended neurons, IT is of fundamental importance to distribute cargo over hundreds of micrometers and is likely finely tuned and balanced to control receptor distribution. Accordingly, IT of neosynthetized AMPAR plays a crucial role to transport them down the dendrites from the Golgi apparatus or Golgi outposts where they are matured after synthesis in the ER. After being released from the Golgi, the secretory vesicles containing the newly-synthesized AMPAR are trafficked to the PM through interaction with adaptor proteins and molecular motors to be finally exocytosed at the PM (Schwenk, Boudkkazi et al., 2019). We and others provided evidence that AMPAR IT is highly modulated by neuronal activity and this suggests that regulation of IT might be a core constituent of the control of synaptic strength during various forms of synaptic plasticity in neuron (Hangen et al., 2018, Hoerndli, Wang et al., 2015). However, despite its potential key role in synaptic regulation, and probable involvement in synaptopathies such as Huntington or Alzheimer diseases (Mandal, Wei et al., 2011), the molecular mechanism that are involved in regulation of AMPAR IT nevertheless remain largely unknown.

AMPAR IT is difficult to study in vertebrates due to the lack of reliable labelling methods and current limitations of imaging systems for detecting fast-moving, low contrast small vesicles. In cultured rat hippocampal neurons, we have overcome these two hurdles using (1) a molecular tool allowing the retention and on-demand release of the newly synthesized AMPAR from the ER/Golgi and (2) the photobleaching of a portion of a dendrite followed by fast video acquisition (Hangen et al.,2018, Rivera, Wang et al., 2000). This allowed the characterization of GluA1 AMPAR subunit IT, that form homomeric calcium-permeable receptors, and can be inserted at synapses during synaptic plasticity (Plant, Pelkey et al., 2006, Sanderson, Gorski et al., 2012).

We found that during chemically induced Long Term Potentiation of synaptic transmission (cLTP), the number and velocities of GluA1 containing vesicles are increased compared to the basal state. These changes in vesicles velocities may be due to the diversity of molecular motors associated with AMPAR, although the exact motors involved are unknown (Hanley, 2014). Molecular motors associate with their cargo through intermediates components, such as adaptor, scaffold and transmembrane proteins (Klopfenstein, Vale et al., 2000). AMPAR are part of a macromolecular complex composed of the receptor *per se* surrounded by a set of associated auxiliary (Bissen, Foss et al., 2019) and cytosolic proteins. Some of these intracellular partners have been shown to be associated with motors proteins and can modulate AMPAR surface expression.

Of particular interest in this regard, 4.1N and SAP97 intracellular proteins are directly and specifically associated with GluA1 C-terminal (C-ter.) domain, the most variable domain between the different AMPAR subunits (Diering & Huganir, 2018, Sans, Racca et al., 2001, Shen, Liang et al., 2000). The C-ter. domain of GluA1 is particularly interesting for the regulation of IT as mutations on this domain modulate its transport, and could be responsible for its upregulation during cLTP (Hangen et al., 2018). However, in the recent years, the C-ter. domain has been under intense scrutiny and its role in mediating synaptic plasticity has been debated. On the one hand, the C-ter. domain of native GluA1 and GluA2 has been suggested to be necessary and sufficient to drive NMDA receptor dependent LTP and LTD respectively (Zhou, Liu et al., 2018). On the other hand, expression of heteromeric receptor containing the GluA1 subunit lacking the C-ter. domain maintains a normal basal trafficking and LTP at CA1 synapses in acute hippocampal slices (Diaz-Alonso, Morishita et al., 2020). The method we developed (Hangen et al., 2018) and results reported here will fuel this debate and allow to determine the exact contribution of GluA1 C-ter. domain for the regulation of IT properties of newly synthetized GluA1 subunit.

In red blood cells, the protein 4.1 (4.1R) is critical for the organization and maintenance of the spectrin–actin cytoskeleton and for the attachment of the cytoskeleton to the cell membrane. 4.1N, the neuronal form of 4.1, may function to confer stability and plasticity to the neuronal membrane via interactions with multiple binding partners such as spectrin–actin-based cytoskeleton, integral membrane channels and receptors. In neurons, 4.1N associates specifically with GluA1 and colocalizes with AMPAR at excitatory synapses (Walensky, Blackshaw et al., 1999). The C-ter. domain of 4.1N mediates the interaction with the membrane proximal region of GluA1. It has been suggested that 4.1N regulates AMPAR trafficking by providing a critical link between the actin cytoskeleton and AMPAR (Shen et al., 2000). Phosphorylation of S_816_ and S_818_ residues in GluA1 regulates activity dependent GluA1 insertion at the PM by enhancing the interaction between 4.1N and GluA1. This suggests that 4.1N is important for the expression of LTP, but doesn’t affect basal synaptic transmission (Lin, Makino et al., 2009). However, while the regulation of GluA1 exocytosis by binding to 4.1N has been established, its potential involvement in AMPAR IT still remains unknown.

SAP97, another important GluA1 C-ter. domain interactor, is a member of the MAGUK family of proteins that play a major role in the trafficking and targeting of membrane ion channels and cytosolic structural proteins in multiple cell types (Fourie, Li et al., 2014). Within neurons, SAP97 is localized throughout the secretory trafficking pathway and at the postsynaptic density (PSD). The role of SAP97 in the control of synaptic function is still unclear despite the fact that the PDZ2 domain of SAP97 interacts directly with the last 4 amino acids of GluA1 (Cai, Coleman et al., 2002). The interaction between SAP97 and GluA1 occurs early in the secretory pathway, while the receptors are in the ER or cis-Golgi, and participates in its forward trafficking from the Golgi to the PM suggesting that SAP97 acts on GluA1 solely before its synaptic insertion and that it does not play a major role in anchoring AMPAR at synapses (Fourie et al., 2014, Sans et al., 2001). SAP97 is a protein known for its involvement in NMDAR (Jeyifous, Waites et al., 2009) and AMPAR IT, thanks to its role as an adaptor protein between GluA1 and the actin based motor MyoVI (Wu, Nash et al., 2002). However, the role of SAP97 in the trafficking and synaptic localization of AMPAR is still debated with conflicting results reported (Fourie et al., 2014, Kay, Tsan et al., 2022, Zhou, Zhang et al., 2008). Moreover, its role in the induction and maintenance of LTP is yet not well characterized.

Here, we have a unique experimental pipeline that allows us to differentiate IT from exocytosis of a given protein by measuring them independently. We report the role of the interactions between 4.1N and SAP97 with the C-ter. domain of GluA1 by analyzing IT and exocytosis of newly synthetized GluA1 deleted for this domain under basal conditions and during synaptic activity. We identify different roles of the interaction between 4.1N-GluA1 and SAP97-GluA1 during basal transmission and after induction of cLTP in hippocampal cultured neurons.

## Materiel and methods

### Molecular Biology

cDNAs of interest are cloned in the ARIAD system (Hangen et al., 2018) then in the Tet-on vector for cLTP experiments. Mutations and deletions are performed by directed mutagenesis and controlled by sequencing before use. sh-RNA against 4.1N are a gift from Huganir’s lab (Lin et al., 2009). Sh-RNA against SAP97 (sequence: GATATCCAGGAGCATAAAT) was clone in a FHUG vector expressing also GFP

### Cell culture

Sprague–Dawley pregnant rats (Janvier Labs) were killed according to the European Directive rules (2010/63/EU). Dissociated hippocampal neurons from E18 embryos were prepared as described previously (Kaech & Banker, 2006) at a density of 300,000 cells per 60-mm dish on poly-L-lysine pre-coated coverslips. Neurons were transfected with the cDNA using Ca^2+^ method at 8-11 days in vitro. Proteins of interest are expressed for 3 to 6 days depending of the experiments. For cLTP experiments expression is started two days before the experiments by addition of 500 nM of doxycycline in the media. All experiments were performed in accordance with the European guidelines for the care and use of laboratory animals, and the guidelines issued by the University of Bordeaux animal experimental committee (CE50; Animal facilities authorizations A3306940, A33063941).

COS-7 cells were maintained in Dulbecco’s Modified Eagle’s Medium (DMEM 4,5G/L + GLUT & PYRUVATE 500, Eurobio) with 10% fetal bovine serum (Eurobio) and 1% L-glutamine (Gibco). Transfection was done with the Xtreme gene HP DNA transfection kit (Roche) following the manufacturer protocol.

### Immunoprecipitation

All subsequent steps were performed at 4 °C. Cells are solubilized in a lysis buffer all (in mM: 50 Hepes pH7.3, 0.5 EDTA, 4 EGTA, 150 NaCl, 1% Triton-X100 and 10μg/mL antiproteases: Leupeptin, Pepstatin, Aprotinin, Pefabloc, Mg132) and centrifuged for 15 min. at 15000G. Supernatant is cleared on the resin for one hour and incubated with the antibodies of interest during 2 hours followed by incubation with protein-A Sepharose overnight. Beads are rinsed with lysis buffer and lysis buffer containing 500 mMol of NaCl. Beads are eluted with Laemmli sample buffer. Western blots are revealed with an Odyssey CLX machine (LI-Cor) by fluorescence method.

### Immunocytochemistry

After addition of 1 μMol of D/D Solubilizer (Ariad Ligand, AL) in the cell culture medium, neurons were incubated for different time. Ten minutes before fixation extracellular labeling of tagged receptors was performed with monoclonal anti-GFP (1/1000) or polyclonal anti-DSRed (1/1000) antibodies incubated in the media at 37°C. Neurons are then fixed with 4% paraformaldehyde, 4% sucrose, rinsed with 50 mM of NH4Cl and transferred in PBS/BSA (1%). Secondary antibody anti-mouse Alexa-fluo588 or anti-rabbit Alexa-fluo488 were applied for 20 minutes at room temperature. Coverslips were mounted on glass slides in Fluoromount-G media (Lonza, ref: 0100-01). Images were collected on an upright Leica DM5000 epifluorescence microscopy with LED light source using a 40x oil objective.

### Quantification

For quantification of extracellular labeling of GFP-GluA1, surfaces of interest were drawn by hand following the total GFP labeling. Images were processed using ImageJ software (Rasband, W.S., ImageJ, U. S. NIH, Bethesda, Maryland, USA, http://imagej.nih.gov/ij/, 1997-2012). Briefly, quantifications were performed by first asking the user to draw sets of regions of interest: one in the background (devoid of any structure), one in the proximal and one in the distal dendritic shaft. For each of them, average intensity and area were retrieved. Results are expressed as the mean intensity fluorescence at the PM. Statistics were performed on Prism software using unpaired t test. Value are in mean +/− SEM. Asterisks notify the following significance levels: p < 0.05 (*), p < 0.01 (**), p < 0.001 (***) and p < 0.0001 (****).

### Videomicroscopy

The videos were acquired on an inverted Leica DMI6000B, equipped with a spinning-disk confocal system (Yokogawa CSU-X1, beam lines: 491 nm, 561 nm), EMCCD camera (Photometrics Quantem 512), and a HCX PL Apo 100X 1.4 NA oil immersion objective. The microscope was driven by the Metamorph software (Molecular Devices, Sunnyvale, USA) and the acquisition took place at 37°C using a Life Imaging Services chamber. The coverslips were mounted in a Ludin chamber filled with 1ml of tyrode media of appropriate osmolarity (15mM glucose, 100mM NaCl, 5mM KCl, 2mM MgCl2, 2mM CaCl2, 10mM HEPES, 247 mosm/l) onto an inverted microscope. 1 μM of AL was added in the tyrode to release the constructs from the ER.

For the basal experiments, during the first 30 minutes, the positions of the expressing cells were saved under the 40X objective. Videos were acquired as follows in the red channel with the 100X objective during the next 30 minutes: during the first 10 seconds, 10 images of 100ms exposure were acquired, followed by the photobleaching of a portion of a dendritic shaft (~60μm^2^, 561nm laser) followed by a one-minute stream of 600 frames of 100 msec exposure to image the transport of the vesicles. For the sh experiments an image was acquired in the green channel prior to the video to confirm the co-transfection of the cell. For each coverslip 4-10 cells were imaged.

For cLTP experiments the positions were saved during the first 20 minutes in a tyrode media (25mM Glucose, 20mM HEPES, 150mM NaCl, 3.5mM KCl, 2 mM MgCl2, 2mM CaCl2, 304 mosm/l). The tyrode was then replaced by a magnesium free tyrode (25mM glucose, 20mM HEPES, 150mM NaCl, 3.5mM KCl, 2mMCaCl2, 200μM glycine, 20 μm bicuculline, 300 mosm/l) for 4 minutes to induce cLTP. Then original tyrode was put back in the Ludin. The videos were acquired 25 minutes after cLTP induction, using the same acquisition parameters as for basal. Cells having less than 5 vesicles / 20 μm^2^/ min. were excluded for the IT analysis of the GluA1 WT condition.

Analyses of the videos are performed using ImageJ program with a help of “Kymo-Tool-Set” software homemade (Hangen et al., 2018). Statistics were performed on Prism software using unpaired t test. Value are in mean +/− SEM. Asterisks notify the following significance levels: p < 0.05 (*), p < 0.01 (**), p < 0.001 (***) and p < 0.0001 (****).

### Materials and antibodies

Ariad ligand (AL), D/D Solubilizer from Takara, ref: 635054. Anti-GFP from Sigma Aldrich (ref: 11814460001), anti-Ds-Red from Ozyme (ref: 632496), anti-myc from Merck Millipore (ref: 6549), anti-4.1N from BD Biosciences (ref: 611836) anti SAP97 from NeuroMab (ref:75-030), anti-GluA1 from NeuroMab (ref: 75-327). Secondary antibodies from Molecular Probes: goat anti-mouse Alexa Fluor-568 (ref: A11004), goat anti-rabbit Alexa Fluor-488 (ref: A11008); from Li-Cor: goat anti-mouse (ref: 926-32210), goat anti-rabbit (ref: 926-68021).

## Results

To study the properties of AMPAR IT, we used cDNA constructs to express GluA1 and its different mutants subcloned in the Ariad system and tagged at its N-terminus (ARIAD-Tag-GluA1) (Hangen et al.,2018). With this technology, receptors are retained in the ER in the basal state thanks to a conditional aggregation domain. Receptor release from the ER and follow-up secretion is tightly controlled with a cell-permeant drug (D/D solubilizer or Ariad ligand: AL) that disrupts aggregation (Rivera et al., 2000). The synchronized release of receptors triggered by addition of AL allows expressed proteins to progress through the secretory pathway in a synchronous manner, particularly adapted to monitor IT. Important features of this system include (1) no or low basal secretion and (2) a rapid and high level of secretion in response to the addition of AL (Hangen et al., 2018). This allowed us to measure three main parameters of AMPAR intracellular trafficking. First, the total number of GluA1 containing vesicles after synchronized release of receptors was used as a measure to ER/Golgi export efficiency. Second, GluA1 vesicle transport properties (speed, fraction of time spent moving or pausing) were measured. Third, we determined the kinetic and extent of GluA1 appearance on the cell surface by live immunolabeling at various times after release. The comparative measurement of these different parameters allowed us to decipher finely the regulatory steps of GluA1 intracellular transport.

4.1N and SAP97 are important proteins implicated in the regulation of AMPAR PM localization. Among all AMPAR subunits, these two proteins are specifically associated with GluA1 C-ter. domain (Lin et al., 2009, Rouach, Byrd et al., 2005, Schwenk, Baehrens et al., 2014), (Sup. Fig. 1A). We analyzed how interactions between these associated proteins and GluA1 regulate AMPAR IT in basal conditions and after induction of chemical long term potentiation (cLTP) in cultured rat hippocampal neurons.

### GluA1 intracellular transport and exocytosis without 4.1N or SAP97

4.1N and SAP97 participate to the biosynthesis and processing of AMPAR in the hippocampus (Sans et al., 2001, Shen et al., 2000). Previous studies established that knocking down 4.1N by expression of a specific sh-RNA substantially reduced GluA1 insertion frequency, indicating that 4.1N is critical for GluA1 insertion at the PM (Lin et al., 2009). On the other hand, SAP97 has been shown to associate with GluA1 containing AMPAR while they are in the ER, with SAP97 dissociating from the receptor at the PM (Sans et al., 2001).

We decided to knock down each of these two proteins independently and analyze IT and externalization of newly synthetized GluA1 in basal condition. We expressed sh-RNA against 4.1N and SAP97 or their corresponding control (scramble) and analyzed the trafficking of ARIAD-Tag-GluA1 after addition of the ligand to release the protein from the ER (Fig. 1).

**Figure1:**
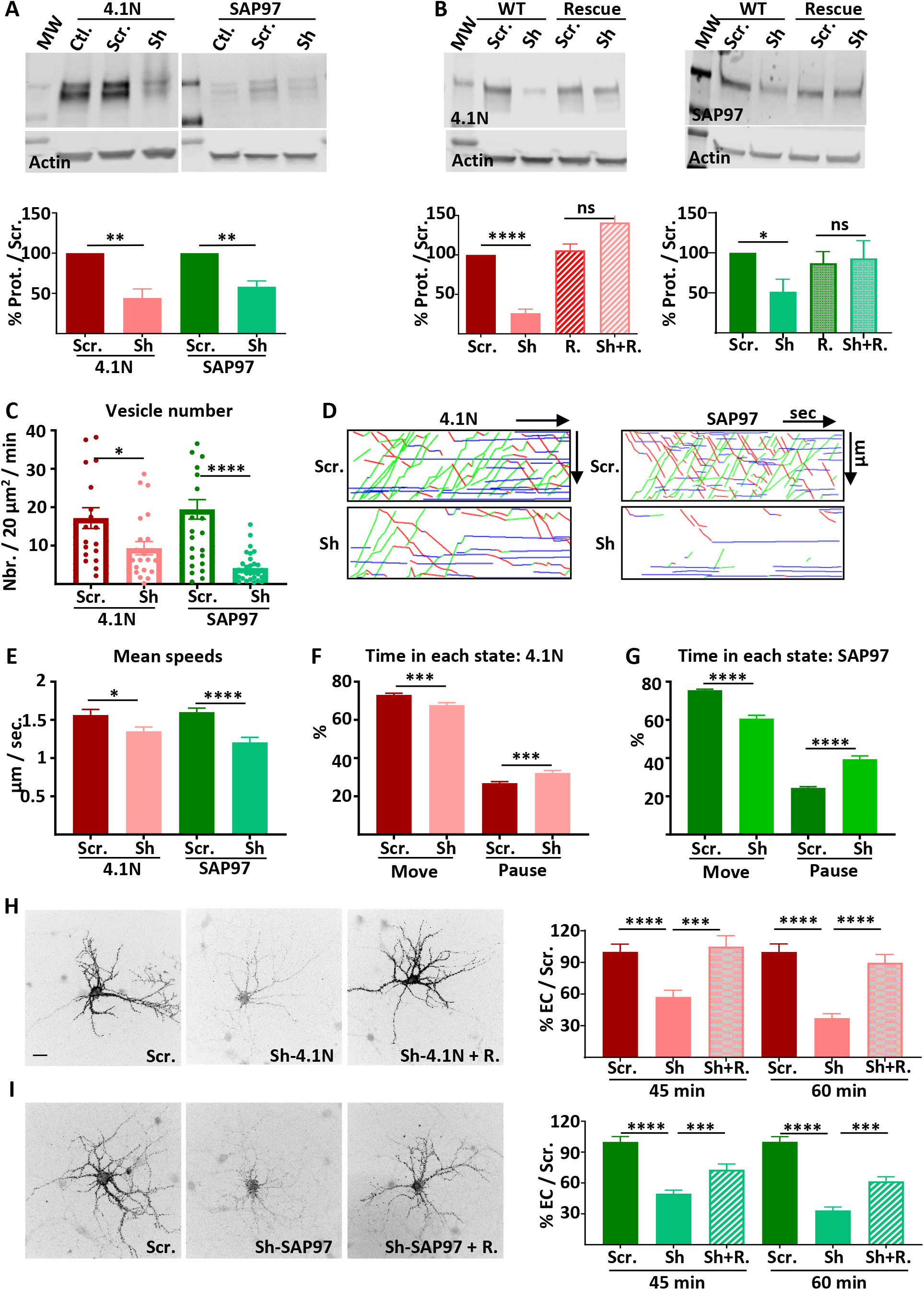
Intracellular transport and exocytosis of GluA1 are dependent of the expression of 4.1N and SAP97. **A-** Top: Western blots of 4.1N and SAP97 expression in cultured rat hippocampal neurons after virus infection with scramble-RNA (scr.) or Sh-RNA against 4.1N and SAP97; bottom: quantification of proteins normalized with actin. **B-** Top: Western blots showing the expression of 4.1N and SAP97 WT and rescue after transfection of the proteins in COS-7 cells and bottom: corresponding quantifications normalized with actin. **C-G-** Parameters of intracellular transport of Ariad-TdTom-GluA1 expressed with scramble-RNA (scr.) or sh-RNA against 4.1N and SAP97. **C-** Vesicle number, **D-** Representative kymographs of the routes of the vesicles in function of the time in video, **E-** Mean speeds of the vesicles in control (expression of scr.) and when 4.1N or SAP97 are decreased (expression of sh), **F-G-** Time spent by a vesicle in a moving state (Move) or in pausing state (Pause). **H-** Representative image of live extracellular labeling of Ariad-GFP-GluA1 after 45 and 60 min. of incubation with AL expressed with sh-RNA for 4.1N with or without the corresponding rescue proteins and quantifications. of incubation with AL. **I-** Representative image of live extracellular labeling of Ariad-GFP-GluA1 after 45 and60 min. of incubation with AL expressed with sh-RNA for SAP97 with or without the corresponding rescue proteins and quantifications. of incubation with AL. Scale bar: 25 μm.

We first controlled the efficacy of the sh-RNA in rat cultured hippocampal neurons by expressing viruses containing respectively scramble RNA (scr.), sh-RNA against 4.1N (sh-4.1N) or against SAP97 (sh-SAP97) (Fig. 1A). Expression of the endogenous proteins were significantly decreased by expression of the corresponding sh (% of scr.; sh-4.1N: 44.4 +/− 11.2, sh-SAP97: 58.3 +/− 7.3). To test the specificity of the sh-RNA, we expressed 4.1N or SAP97 or the corresponding rescue proteins in COS-7 cells together with the scr.-RNA or the sh-RNA and quantified expression of 4.1N and SAP97 by western blots analysis (Fig. 1B). As in neurons, expression of sh-RNA decreased the expression of the corresponding wild type proteins (% of scr.; sh-4.1N: 26.2 +/− 10.1, sh-SAP97: 51.4 +/− 15.6) without affecting the expression of the corresponding rescue proteins (% of scr. on rescue proteins; scr.: 105.9 +/− 7.9, sh-4.1N: 141.2 +/− 3.1, scr.: 87.0 +/− 1.5, sh-SAP97: 93.0 +/− 2.2)., showing the specificity of our sh-RNA.

We then analyzed the parameters of GluA1 IT when 4.1N or SAP97 sh-RNA or corresponding scr.-RNA were expressed (Fig. 1C-G). We expressed GluA1 subcloned in the Ariad vector, induced the transport of the protein by addition of the AL and analyzed the transport 30 to 60 minutes after addition of the AL. In both cases, the total number of vesicles transporting GluA1 was decreased, although less drastically when 4.1N was knocked down than when SAP97 was (vesicles / 20 μm^2^ / min; scr. 4.1N: 17.2 +/− 2.7, sh-4.1N: 9.3 +/− 1.7, scr. SAP97: 19.5 +/− 2.5; sh-SAP97: 4.2 +/− 0.7) (Fig. 1C).

For each cell, we traced the corresponding kymographs and calculated the mean speed of the vesicles (Fig. 1D-E, Sup. Fig. 1B). We found similar values for speed for the OUT (from the cell body to the dendrite) and for the IN (from the dendrite to the cell body) directions (Sup. Fig. 1C). We thus decided to pool the speeds of transport of OUT and IN together. The mean speed of the vesicles was only decreased by 15% by expression of the sh-4.1N compared to its scr.-RNA (μm / sec; scr. 4.1N: 1.56 +/− 0.07, sh-4.1N: 1.35 +/− 0.05) (Fig. 1E, Sup. Fig. 1C). When sh-SAP97 was expressed, the mean speed was decreased by 25% compared to the corresponding scr. (μm / sec; scr. SAP97: 1.60 +/− 0.05, sh-SAP97: 1.20 +/− 0.07). Thanks to the kymographs, we calculated the percentage of time spent in the moving and in the pausing states for each vesicle (Fig. 1F-G, Sup. Fig. 1D-E). The time spent moving was also decreased when the sh-4.1N was expressed (12% less) to the benefit of the pausing time (% Move: scr. 4.1N: 73.13 +/− 0.83, sh-4.1N: 67.77 +/− 1.25; % pause: scr. 4.1N: 26.87 +/− 0.83, sh-4.1N: 32.22 +/− 1.25). Indeed, when expression of SAP97 was decreased, the time spent moving by a vesicle was decreased (20 % less) to the benefit of the time spent in pause (% Move: scr. SAP97: 75.57 +/5.89, sh-SAP97: 60.68 +/− 1.74; % pause: scr. SAP97: 24.43 +/− 5.89, sh-SAP97: 39.32 +/− 1.75).

We then analyzed the kinetics of externalization of GluA1 in the same conditions as for the IT experiments. We performed live extracellular labeling of GluA1 45 and 60 min. after addition of the AL on hippocampal rat cultured neurons (Fig. 1H-I). Expression of sh-4.1N decreased massively the externalization of GluA1 compared to its externalization with the corresponding scr. (% of cle.; 45 min. after AL: 57.36 +/− 6.27, 60 min. after AL: 37.29 +/− 4.16) (Fig. 1H). The 4.1N rescue protein could prevent this decrease in the rate of externalization when expressed together with the sh-4.1N (% of cle.; 45 min. after AL: 105.00 +/− 10.34, 60 min. after AL: 89.73 +/− 7.81). Expression of sh-SAP97 also decreased the rate of externalization of GluA1 to the same extent as sh-4.1N did (% of cle.; 45 min. after AL: 49.64 +/− 3.41, 60 min. after AL: 33.33 +/− 3.29) (Fig. 1I). Expression of SAP97 rescue protein partially restored the externalization of GluA1 (% of cle.; 45 min. after AL: 72.91 +/− 5.58, 60 min. after AL: 61.69 +/− 4.32).

In conclusion, reducing the expression of 4.1N and SAP97 both diminished GluA1 IT in rat cultured hippocampal neurons. However, the effects were all less drastic when 4.1N expression was decreased than when SAP97 was. This was the case for the decrease in the number of vesicles released upon addition of AL, for the decrease in the vesicles speed and for the increase in pausing time. However, we found that the externalization of GluA1 at the PM was equally inhibited by the absence of 4.1N or SAP97. Because down-regulation of 4.1N or SAP97 could have indirect effects on GluA1 transport properties, with then studied the impact of GluA1 mutations that inhibit its interaction with these proteins.

### Regulation of AMPAR traffic by the interaction between GluA1 C-ter. domain and 4.1N or SAP97

GluA1 IT and PM localization are dependent on the expression of 4.1N and SAP97. Knocking down either 4.1N or SAP97 decreases massively the exit of GluA1 from the ER-Golgi and impacts IT and exocytosis of the receptor to the PM. However, the overall impact of SAP97 is more drastic on IT whereas we found the same effect for both conditions on the PM localization of GluA1. This may be because the interaction between 4.1N and GluA1 might be necessary mainly for the exocytosis of the receptor at the PM. We decided to analyze if the interaction between GluA1-4.1N or GluA1-SAP97 is important for the intracellular transport and exocytosis of the newly synthetized receptor in basal synaptic transmission (Fig. 2).

**Figure2:**
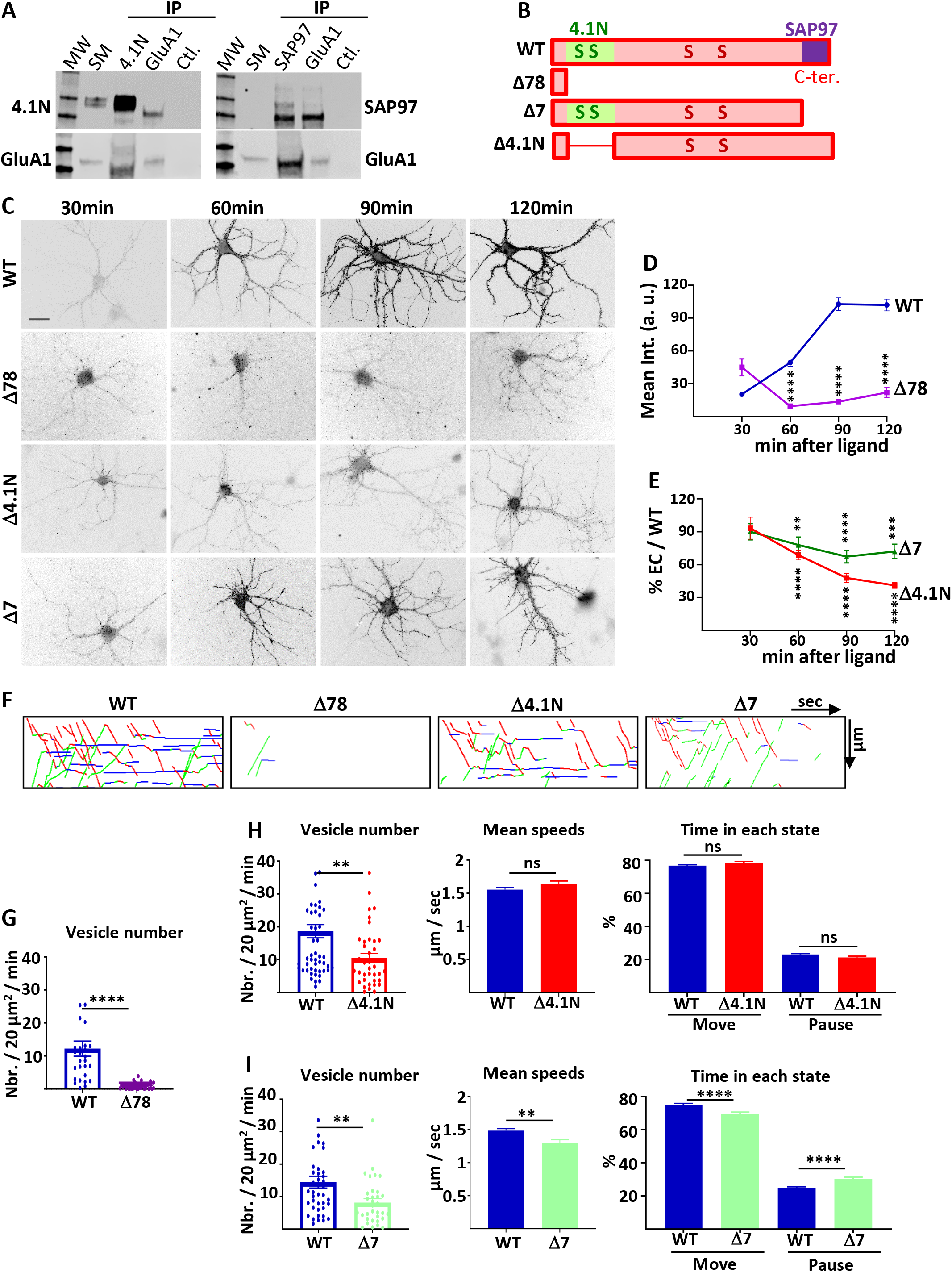
4.1N/GluA1 and SAP97/GluA1 interactions differently regulate GluA1 traffic in basal transmission. **A-** Co-immunoprecipitation of endogenous GluA1 with 4.1N and SAP97 in cultured rat hippocampal neurons. Control (Ctl.) is performed without antibody. Western blot of GluA1, 4.1N and SAP97 as indicated. **B-** Diagram of the different truncated mutants on the C-terminal (C-ter.) domain of GluA1. **C-** Representative images of live labeling of Ariad-GFP-GluA1 after addition of AL during different time as indicated. Scale bar: 25 μm. **D-** Quantification of the exit of Ariad-GFP-GluA1-WT (WT) and Ariad-GFP-GluA1-Δ78 (Δ78) over time after addition of AL. For the WT 100% of exit is taken after 120 min of addition of AL. **E-** Quantification of the exit of Ariad-GFP-GluA1-Δ4.1N (Δ4.1N) and Ariad-GFP-GluA1-Δ7 (Δ7) over time after addition of AL. The 100% values correspond to the value of the WT for the same time of induction. **F-** Traced kymographs for the different mutants. **G-** Number of vesicles detected for the ARIAD-TdTom-GluA1-WT (WT) and the ARIAD-TdTom-GluA1-Δ78 (Δ78). **H-I-** Parameters of intracellular transport: vesicle number, mean speeds and percentage of time in each state for the ARIAD-TdTom-GluA1-Δ4.1N (Δ4.1) (**H**) and ARIAD-TdTom-GluA1-Δ7 (Δ7) (**I**) taking as a 100 % the corresponding values for ARIAD-TdTom-GluA1-WT protein from the same experiments.

We first checked by co-immunoprecipitation experiments if endogenous GluA1 and 4.1N and GluA1 and SAP97 are interacting in our model of rat cultured hippocampal neurons (Fig. 2A). Indeed, we found that immunoprecipitation of 4.1N or SAP97 co-immunoprecipitated GluA1 and that immunoprecipitation of GluA1 co-immunoprecipitated 4.1N and SAP97. It has been shown that GluA1 binds SAP97 by its last 4 amino acids (Leonard, Davare et al., 1998) whereas it binds 4.1N on a peptide domain localized just after its fourth transmembrane domain (Shen et al., 2000). We designed different GluA1 mutants in order to study the impact of these interactions on GluA1 IT (Fig. 2B). We deleted the entire C-ter. domain of GluA1 (deletion of the last 78 amino acids of GluA1 leaving only 4 amino acids after the last transmembrane domain, Δ78) or each of the interaction sites (Δ4.1N for 4.1N and Δ7 for SAP97). For each mutant, we studied their PM localization as a function of the time of incubation with the AL and the characteristic of their IT.

The externalization of newly synthetized receptors expressing GFP-GluA1-Δ78 or GFP-GluA1-WT was monitored by live immunolabeling with an antibody directed against GFP (Fig. 2C, D). Quantification of the GFP staining revealed an almost total disappearance of GluA1-Δ78 externalization (WT versus Δ78, arbitrary unit: 30 min, 20.43 +/− 2.14 versus 45.02 +/− 7.75; 60 min, 49.39 +/− 3.42 versus 9.54 +/− 1.95; 90 min, 102.68 +/− 5.83 versus 13.64 +/− 1.70; 120 min, 101.95 +/− 5.37 versus 22.02 +/− 4.59). This experiment demonstrates that the C-ter. domain is necessary for newly synthetized GluA1 to be externalized at the PM, even two hours after triggering GluA1 ER exit.

We then studied if the interaction between GluA1 and 4.1N or SAP97 is necessary for the localization of GluA1 at the PM (Fig. 2C, E, Sup. Fig. 2A). We performed extracellular labeling of GFP-GluA1-WT and the mutants, GFP-GluA1-Δ4.1N and GFP-GluA1-Δ7 respectively deleted for their binding site for 4.1N or SAP97, after induction of IT by addition of the AL during different times. For analysis of these experiments, we normalized the externalization values of the mutants to that of GFP-GluA1-WT at the corresponding times of incubation with AL in paired experiments. Both mutants were less exocytosed than GluA1-WT after 30 min to 2 hours of addition of the AL. At each time point, the exocytosis of GluA1 lacking the 4.1N binding site was more decreased than that of GluA1 lacking the SAP97 site (Δ4.1N: 30 min, 93.22 +/− 10.00; 60 min, 68.75 +/− 4.43; 90 min, 47.94 +/− 4.01; 120 min, 41.19 +/− 2.70. Δ7: 30 min, 90.07 +/− 7.42; 60 min, 77.98 +/− 7.26; 90 min, 67.35 +/− 5.68; 120 min, 72.04 +/− 6.60). When quantified at 90 and 120 min. the difference between the two mutants was highly significant (Sup. Fig. 2A). This result shows that interaction of GluA1 with 4.1N or SAP97 is important for surface expression of newly synthetized GluA1. The extracellular labeling of GluA1 is less important when the site of interaction of 4.1N is deleted than when the site of interaction of SAP97 is deleted.

This lack of normal exocytosis of the mutants can be due either to an inhibition of externalization or to a decrease of their IT. We thus analyzed the IT parameters of the different mutants taking GluA1-WT as a control (Fig. 2F-I, Sup. Fig. 2B, C). For these experiments, we expressed ARIAD-TdTom-GluA1, WT or mutants, in order to be in the best conditions to detect the transport vesicles. We first analyzed the IT of the Δ78 mutant (Fig. 2F-G). The number of vesicles transporting this mutant was very low compared to GluA1-WT (vesicles / 20 μm^2^ / min; GluA1-WT: 12 +/− 2.3, Δ78: 0.86 +/− 0.2) and this prevented the analysis of their IT parameters. The C-ter. domain of GluA1 is thus mandatory for the exit of newly synthetized GluA1 from the ER and the Golgi, likely explaining its requirement for GluA1 surface expression.

We then characterized the IT of the mutant deleted for the 4.1N binding site (Δ4.1N) (Lin et al.,2009) (Fig. 2F, H, Sup. Fig. 2B, C). The number of vesicles transporting the protein was decreased compared to GluA1-WT (vesicles / 20 μm^2^ / min; GluA1-WT: 18.68 +/− 2.05, Δ4.1N: 10.52 +/− 1.40). This is in accordance with a decrease of the number of vesicles that we found when 4.1N was knocked down by expression of the sh-4.1N. In contrast, the mean speeds of the vesicles were the same for GluA1-WT and GluA1-Δ4.1N (μm / sec; WT: 1.55 +/− 0.03, Δ4.1N: 1.64 +/− 0.05). The time spent in each state was similar for the two proteins (% Move WT: 76.75 +/− 0.53 %, Δ4.1N: 78.52 +/− 0.78 %; % pause: WT: 23.10 +/− 0.53 %, Δ4.1N: 21.31 +/− 0.78 %). These results indicate that binding of GluA1 to 4.1N is important for its ER or Golgi export and exocytosis at the PM but does not affect its IT once the vesicles are released from the Golgi apparatus.

We then analyzed the IT of the mutant deleted for the last 7 amino acids corresponding to the binding site of SAP97 (Δ7) (Zhou et al., 2008) (Fig. 2F, I, Sup. Fig. 2B, C). For this mutant, the number of vesicles released was decreased compared to GluA1-WT, as for GluA1-Δ4.1N (vesicles / 20 μm^2^ / min; GluA1-WT: 14.45 +/− 1.82 vesicles / 20 μm^2^ / min, Δ7: 8.16 +/− 1.21). We also found a highly significant effect of the Δ7 mutant deletion of the PDZ binding domain on all the GluA1 IT parameters. First, the mean speed was decreased by 12% for the Δ7 mutant (μm / sec; WT: 1.48 +/− 0.03, Δ7: 1.30 +/− 0.05). Moreover, the time spent in a moving state was decreased by 7% and, conversely, the percentage of time in pause was increased by 22 % compared to the WT (% Move: WT: 75.18 +/− 0.70 %, Δ7: 69.67 +/− 1.04 %; % pause: WT: 24.82 +/− 0.69 %, Δ7: 30.33+/− 1.04 %). All these changes, although relatively modest, are significant and can have important functional impacts when cumulated over time. This corresponds to what we found for IT properties when the expression of SAP97 was decreased: modified ER/Golgi export and PM exocytosis, reduced speed and increased time in pause.

### Regulation of AMPAR traffic by the interaction between point mutated GluA1 C-ter. domain with 4.1N

Since we had an impact on GluA1 IT when 4.1N was knocked down, we were surprised by the absence of impact of the Δ4.1N deletion on GluA1 IT. We thus decided to analyze the characteristics of IT with a GluA1 mutant that does not bind 4.1N and has the same C-ter. domain length (Fig. 3, Sup. Fig. 3). During LTP, protein kinase C (PKC) phosphorylates the serine 816 (S_816_) and serine 818 (S_818_) residues in the GluA1 C-ter. domain. These phosphorylations enhance 4.1N binding to GluA1 and facilitates GluA1 insertion at the PM. When these two serines are replaced by alanines, the interaction with 4.1N is abolished (Lin et al., 2009). We constructed the corresponding S_816_A-S_818_A (AA) or the LTP like constitutively phosphorylated S_816_D-S_818_D (DD) mutants of ARIAD-TdTom-GluA1 and ARIAD-GFP-GluA1 (Fig. 3A). These constructs allowed us to study with more specificity the impact of the interaction between 4.1N and GluA1.

**Figure3:**
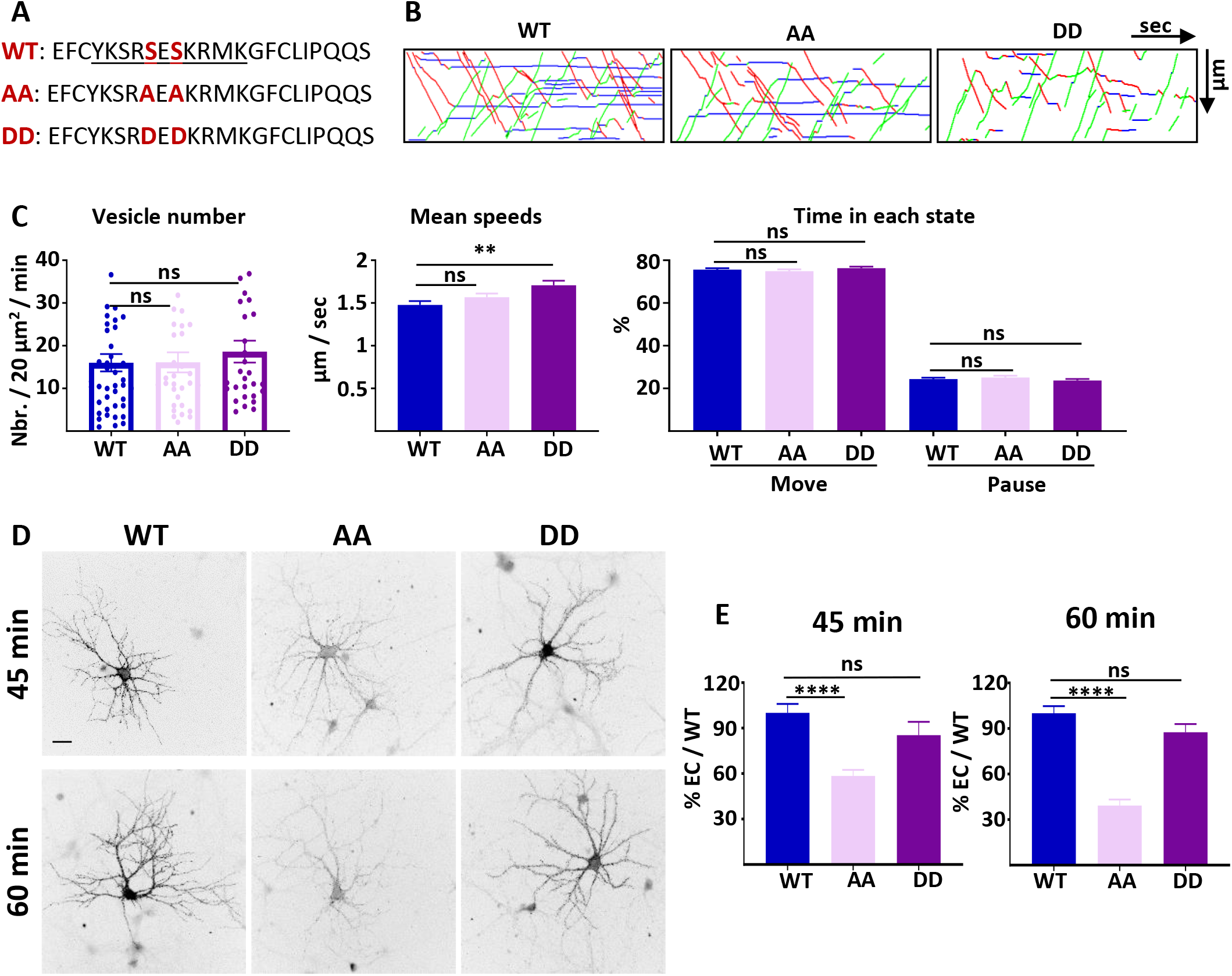
4.1N/GluA1 interaction is only necessary for the exocytosis of GluA1 in basal condition. **A-** Amino acid sequences representing the binding site of 4.1N on the C-ter. domain of GluA1 and mutations corresponding to the S_816_A S_818_A (AA) and S_816_D S_818_D (DD) mutants. **B-** Representative traced kymographs for ARIAD-TdTom-GluA1-WT (WT), ARIAD-TdTom-GluA1-AA (AA) and ARIAD-TdTom-GluA1-DD (DD). **C-** Parameters of intracellular transport: vesicle number, mean speeds and percentage of time in each state for the WT, AA and DD. **D-** Representative images of live labeling of Ariad-GFP-GluA1-WT (WT), Ariad-GFP-GluA1-AA (AA) and Ariad-GFP-GluA1-DD (DD) after addition of AL during different time as indicated. Scale bar: 25 μm. **E-** Quantification of live labeling after 45 and 60 min. of incubation with AL normalized on the WT levels after the same time of incubation with AL.

We first performed IT experiments with these GluA1 mutants (Fig. 3B, C; Sup. Fig. 3). The number of vesicles was similar for the WT AA and DD mutants (vesicles / 20 μm^2^ / min; GluA1-WT: 16.01 +/− 2.05, AA: 16.10 +/− 2.34, DD: 18.61 +/− 2.56) (Fig. 3B, C; Sup. Fig. 3A). We then analyzed the mean speed of the vesicles containing each protein (μm / sec; WT: 1.48 +/− 0.04, AA: 1.57 +/− 0.04, DD: 1.71 +/− 0.05) (Fig. 3C, Sup. Fig. 3B, C). As for the Δ4.1N mutant, we did not find any difference in the mean IT speed between WT and AA proteins. However, the mean speed for the DD mutant was significantly increased compared to the WT, specifically for the OUT direction (Sup. Fig. 3B). The frequency distribution of the speeds was identical for the three conditions in both OUT and IN directions (Sup. Fig. 3C). The percentage of time spent in moving and pausing states was also identical for the three proteins WT, AA and DD (Move: WT: 75.64 +/− 0.69 %, AA: 74.90 +/− 0.84 %, DD: 76.31 +/− 0.75 %; Pause: WT: 24.36 +/− 0.69 %, AA: 25.10 +/− 0.84 %, DD: 23.70 +/− 0.75 %) (Fig. 3D, Sup. Fig. 3D). This shows that, in the basal state, GluA1 IT is not dependent on the binding of 4.1N since the AA mutant has the same properties than the WT. This result is in accordance with the observation reported above that the GluA1-WT and GluA1-Δ4.1N IT are similar. We thus conclude that the impact of knocking down 4.1 N on GluA1 IT, in basal condition, does not originate from an interaction between GluA1 and 4.1N.

We then measured the export to the PM of GluA1 WT and the corresponding mutants 45 min and 60 min after addition of AL (Fig. 3D, E). Already after 45 min. of IT induction, we found that the exit of GluA1 was strongly decreased for the GluA1-AA mutant but comparable with the WT for the DD mutant (% of the WT; AA: 58.44 +/− 3.99; DD: 85.39 +/− 8.75). This difference in exocytosis of GluA1 was accentuated when the neurons were incubated during 60 min with the AL (% of the WT; AA: 39.27 +/− 4.01; DD: 85.57 +/− 5.39).

In conclusion, during basal transmission, the interaction between GluA1 and 4.1N has a fundamental role for the ER/Golgi export and for the exocytosis at the PM of GluA1 without regulating its IT.

### Role of SAP97 on the regulation of GluA1 traffic during cLTP

We have shown previously that GluA1 IT is strongly increased 25-40 minutes after induction of cLTP in cultured rat hippocampal neurons (Hangen et al., 2018). In this condition, the number of vesicles is increased by 140%, the speed is higher and the pausing time is lower. Here, we applied the same cLTP protocol to investigate the role of SAP97 (and then 4.1N below) in the regulation of the late phase of cLTP (200 μM Glycine, 20 μM bicucullin, 0 mM Mg^++^ during 3 min. and then return to normal media). Videos were acquired 25-40 minutes after induction of cLTP. For some experiments, a video was acquired before induction for 1-2 cells expressing GluA1-WT control to verify the efficacy of our cLTP protocol (Sup. Fig. 4A). We analyzed the ER/Golgi export, transport and plasma membrane export properties of GluA1 after cLTP in function of both the level of expression of SAP97 and the binding of GluA1 to SAP97 by expressing scramble or sh-SAP97 and comparing IT between GluA1-WT and GluA1-Δ7 mutant (Fig. 4, Sup. Fig. 4).

**Figure 4:**
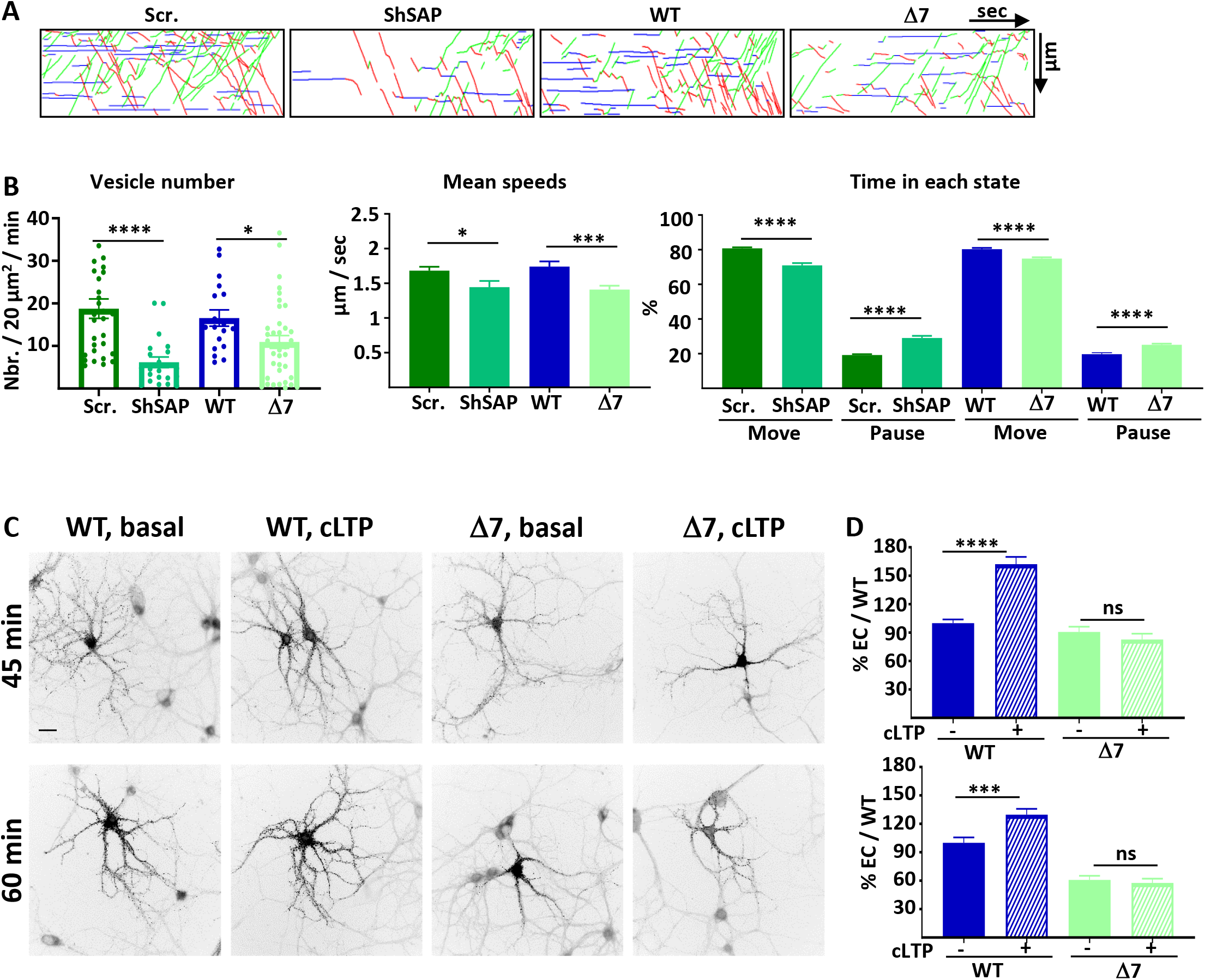
SAP97 and SAP97/GluA1 interaction regulate GluA1 trafficking during cLTP. **A-** Representative traced kymographs for ARIAD-TdTom-GluA1-WT co-transfected with scramble-RNA (Scr.) or with sh-SAP97 (ShSAP), ARIAD-TdTom-GluA1-WT (WT) and ARIAD-TdTom-GluA1-Δ7 (Δ7). **B** IT parameters of GluA1 in the different conditions as indicated 25-40 minutes after induction of cLTP: Vesicle number, mean speeds of the vesicles and time in each state (Move and Pause). **C-** Representative images of neurons for each conditions 45-60 min after addition of the AL, 25-40 after induction of cLTP. Scale bar: 25 μm. **D-** Quantification of live immunolabeling before and after induction of cLTP for the GluA1-WT (WT) and GluA1-Δ7 (Δ7) mutant.

The number of vesicles containing GluA1 after cLTP was strongly decreased by expression of sh-SAP97 as compared to cLTP in control conditions (vesicles / 20 μm^2^ / min; scr.: 18.77 +/− 2.29, Sh-SAP97: 6.18 +/− 1.25) and when we expressed GluA1-Δ7 compared to GluA1-WT (vesicles / 20 μm^2^ / min; WT: 16.55 +/− 1.92, Δ7: 10.95 +/− 1.50) (Fig. 4A, B, Sup. Fig. 4B). These effects were identical to those observed in basal conditions both for Sh-SAP97 and Δ7 mutant conditions (% decrease of vesicles: control/Sh-SAP97 in basal: 78.4 %, after cLTP: 67.1%; control/Δ7 in basal: 27.2%, after cLTP: 33.8 %). Similarly to the effects observed in basal conditions, the mean speeds were also decreased after cLTP by both sh-SAP97 and GluA1-Δ7 truncation (μm / sec; scr.: 1.68 +/− 0.06, Sh-SAP97: 1.45 +/− 0.09) and (μm / sec; WT: 1.74 +/− 0.08; Δ7: 1.41 +/− 0.06) (Fig. 4B, Sup. Fig. 4C). The time spent in the moving state were also similarly decreased compared to the corresponding controls (Move: scr.: 80.82 +/− 0.53 %, Sh-SAP97: 70.99 +/− 1.29 %; pause: scr.: 19.18 +/− 0.53 %, Sh-SAP97: 29.00 +/− 1.30 %) and (Move: WT: 79.63 +/− 0.75 %, Δ7: 76.45 +/− 0.73 %; pause: WT: 20.37 +/− 0.75 %, Δ7: 23.55 +/− 0.73 %) (Fig. 4B). These effects are in accordance with the impact that we already found during basal transmission.

We next analyzed the impact of the GluA1-Δ7 truncation on its externalization to the PM after induction of cLTP (Fig. 4C, D). We induced cLTP 20 min after addition of AL and performed live immunolabeling of neosynthetized GluA1 25 min (total with AL 45 min) or 40 min (total with AL 60 min) after induction of cLTP. cLTP significantly increased the exit of GluA1-WT (% to the basal; exit 45 min: 162.24 +/− 7.71; 60 min: 129.63 +/− 6.22) but had no impact on the externalization of GluA1-Δ7, neither at 45 min nor at 60 min (%; exit 45 min: basal 90.71 +/− 5.5 % of WT, LTP 82.78 +/− 6.09 %; exit 60 min: basal 60.82 +/− 4.27 %, LTP 57.62 +/− 4.50 %). Altogether, this shows that downregulating SAP97 or preventing its binding to GluA1 abolished the cLTP induced regulation of GluA1 ER/Golgi exit, IT and PM externalization.

### Role of 4.1N on the regulation of GluA1 traffic during cLTP

Disrupting 4.1N-dependent GluA1 PM insertion decreases surface expression of GluA1 and the expression of long-term potentiation showing that 4.1N is required for activity-dependent GluA1 insertion in rodents (Lin et al., 2009). These experiments were performed by directly visualizing individual insertion events of the AMPAR subunit GluA1 at the PM. It was thus particularly interesting to study if the 4.1N/GluA1 interaction contributes to the regulation of GluA1 IT during synaptic plasticity or only to the exocytosis of GluA1. We thus next studied the contribution of 4.1N to cLTP-induced regulation of GluA1 IT. In basal condition, when GluA1 is not associated with 4.1N (GluA1-AA), IT is normal compared to GluA1-WT. However, the interaction between GluA1 and 4.1N is important for the exit of GluA1 from the ER/Golgi and the insertion at the PM. We thus analyzed the transport of GluA1 during cLTP upon downregulation of 4.1N by sh-4.1N and when the interaction between 4.1N and GluA1 is abolished such as for the GluA1-Δ4.1N mutant. As for SAP97, we applied the cLTP protocol in cultured rat hippocampal neurons (Hangen et al., 2018) and compared IT of GluA1 or mutant 25-40 minutes after induction of cLTP (Fig. 5, Sup. Fig. 5).

**Figure 5:**
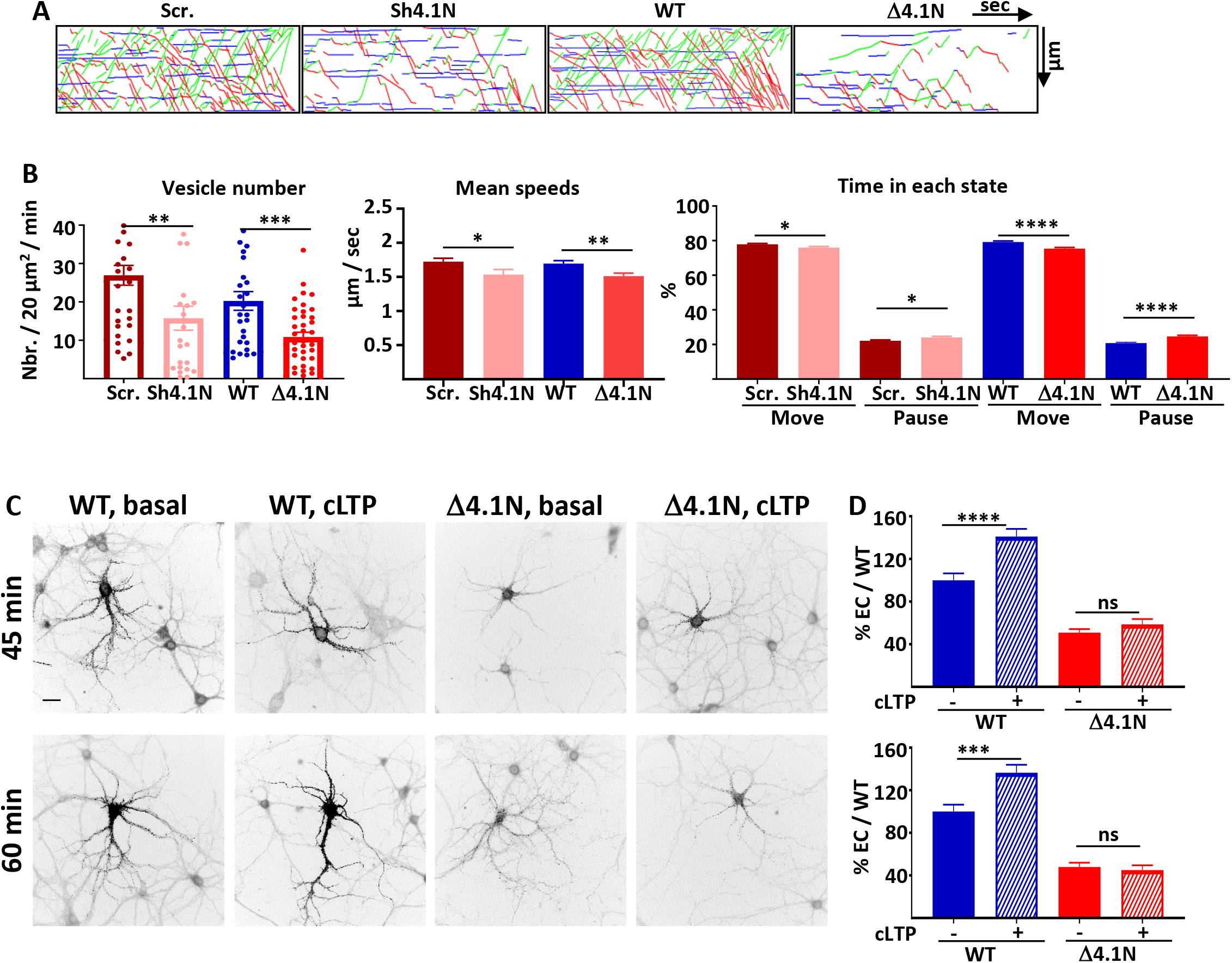
4.1N and 4.1N/GluA1 interaction regulate GluA1 trafficking during cLTP. **A-** Representative traced kymographs for ARIAD-TdTom-GluA1-WT co-transfected with scramble-RNA (Scr.) or with sh-4.1N (Sh4.1N), ARIAD-TdTom-GluA1-WT alone (WT) and ARIAD-TdTom-GluA1-Δ4.1N (Δ4.1N). **B-** IT parameters of GluA1 in the different conditions as indicated 25-40 minutes after induction of cLTP: Vesicle number, mean speeds of the vesicles and time in each state (Move and Pause). **C-** Representative images of neurons for each conditions 45-60 min after addition of the AL, 25-40 min. after induction of cLTP. Scale bar: 25 μm. **D-** Quantification of live immunolabeling before and after induction of cLTP for the GluA1-WT (WT) and GluA1-Δ4.1N (Δ4.1N) mutant.

After induction of cLTP, the number of vesicles was decreased compared to the corresponding controls either by decreasing the expression of 4.1N (vesicles / 20 μm^2^ / min; scr.: 26.94 +/− 2.59, Sh-4.1N: 15.75 +/− 3.13) or by inhibition of the binding by expression of Δ4.1N (vesicles / 20 μm^2^ / min; WT: 20.25 +/− 2.42, Δ4.1N: 10.84 +/− 1.23) (Fig. 5A, B, Sup. Fig. 5A). Without 4.1N the decrease of the number of vesicle transporting GluA1 was comparable after cLTP and in basal condition. In the same way, inhibition of binding of 4.1N on GluA1 decreased similarly IT of GluA1 in basal and cLTP conditions (% decrease of vesicles: control/Sh-4.1N in basal: 45.6 %, after cLTP: 41.5 %; control/Δ4.1N in basal: 43.4%, after cLTP: 46.4 %). The mean speed of IT 25-40 min after induction of cLTP was impacted by expression of the sh-4.1N (μm / sec; scr.: 1.72 +/− 0.05, Sh-4.1N: 1.53 +/− 0.07) (Fig. 5B). It was also decreased when we expressed GluA1-Δ4.1N compare to the speed of GluA1-WT (μm / sec; WT: 1.69 +/− 0.04; Δ4.1N: 1.51 +/− 0.04). This difference of speed translated into a decrease of the moving state to the benefit of pausing time (Move: scr.: 77.81 +/− 0.50 %, Sh-4.1N: 75.95 +/− 0.67 %; pause: scr.: 22.18 +/− 0.50 %, Sh-4.1N: 24.02 +/− 0.67 %) and (Move: WT: 79.23 +/− 0.54 %, Δ4.1N: 75.34 +/− 0.67 %; pause: WT: 20.76 +/− 0.54 %, Δ4.1N: 24.70 +/− 0.67 %) (Fig. 5B).

We then analyzed the exocytosis of newly synthetized GluA1 after induction of cLTP (Fig. 5C, D). The induction of cLTP allows an increase of the externalization of GluA1 WT protein (% of GluA1 basal: 45 min: 141.06 +/− 6.51 %; 60 min: 136.52 +/− 7.47 %). In the same conditions, cLTP had no impact on the externalization at the PM of GluA1-Δ4.1N (% of GluA1 basal: 45 min: 50.80 +/− 3.37 %; 60 min: 45.04 +/− 4.52 %).

Altogether, both suppressing the SAP97 and the 4.1N interaction with GluA1 prevented the cLTP induced regulation of GluA1 ER/Golgi exit, IT and plasma membrane exit.

We found that the parameters of GluA1-Δ4.1N IT are decreased during cLTP. During LTP, protein kinase C (PKC) phosphorylation of the serine 816 (S_816_) and serine 818 (S_818_) residues of GluA1 enhanced 4.1N binding to GluA1 and facilitated GluA1 insertion at the PM (Lin et al., 2009). We thus decided to study the impact of cLTP when GluA1 cannot be phosphorylated at these two sites by using the AA mutant that does not bind 4.1N (Fig. 6, Sup. Fig. 6). This construct allowed us to study directly the impact of the binding of 4.1N after induction of cLTP together with the impact on exocytosis of GluA1 in both conditions.

**Figure 6:**
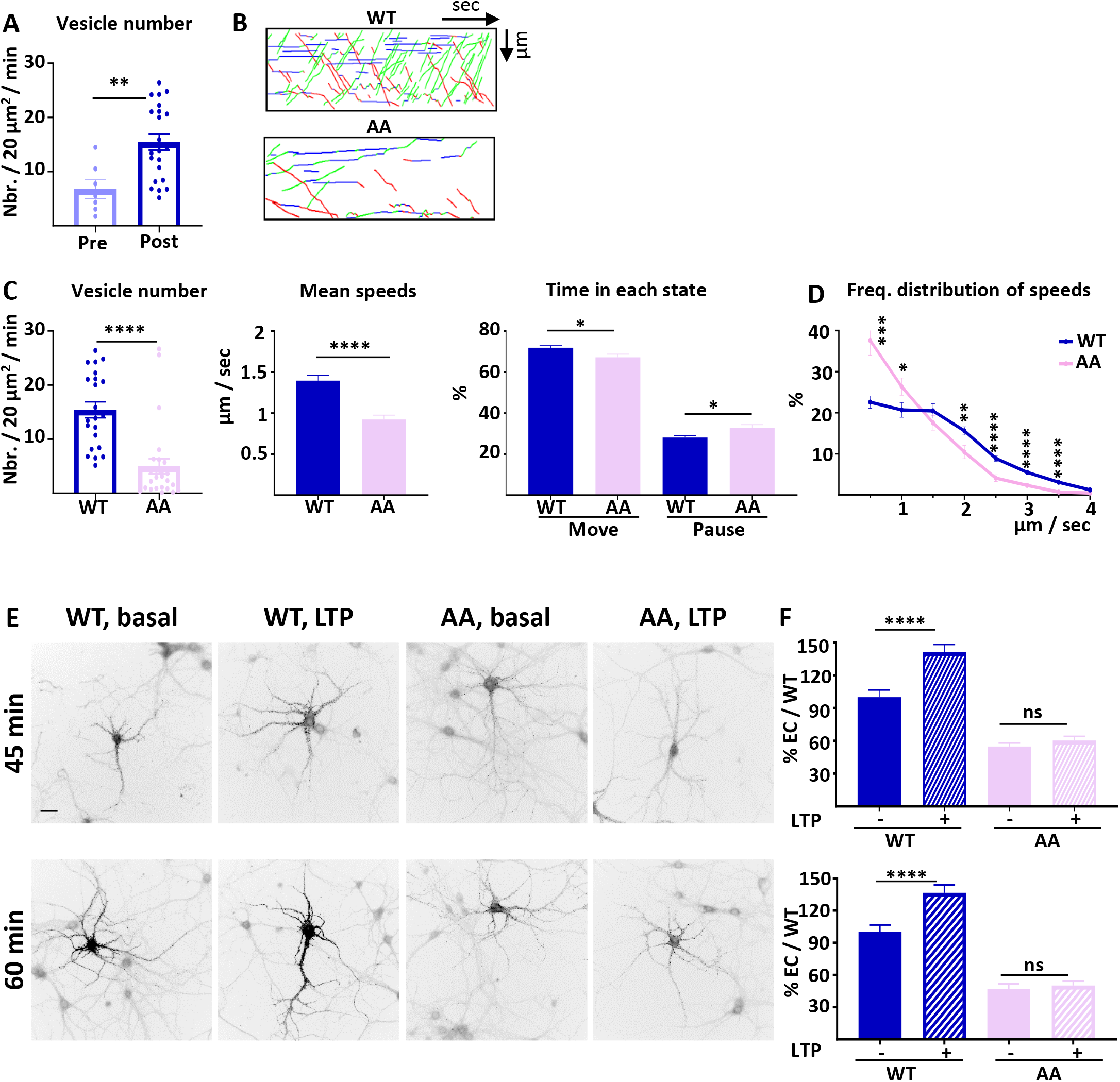
4.1N/GluA1 interaction drives intracellular transport of GluA1 during cLTP. **A-** Number of vesicles before (Pre) and after (Post) induction of cLTP for ARIAD-TdTom-GluA1-WT. **B-** Representative traced kymographs for ARIAD-TdTom-GluA1-WT (WT) and ARIAD-TdTom-GluA1-AA (AA) 25-40 min after induction of cLTP. **C-** IT parameters of GluA1 in the different conditions as indicated 25-40 minutes after induction of cLTP: Vesicle number, mean speeds of the vesicles and time in each state (Move and Pause). **D-** Frequency distribution of speed for the two proteins after induction of cLTP. **E-** Representative images for each conditions 45 min and 60 min after addition of the AL, 25 and 40 min. after induction of cLTP. Scale bar: 25 μm. **F-** Quantification of live labeling of GluA1 in WT and AA conditions, before and after cLTP.

On each set of experiments we verified that the cLTP protocol increased the number of vesicles on a few cells expressing Ariad-TdTom-GluA1 (Hangen et al., 2018) (vesicles / 20 μm^2^ / min; Pre cLTP: 6.77 +/− 1.70, Post cLTP: 13.99 +/− 1.52) (Fig. 6A). After cLTP the number of vesicles for the AA mutant was decreased by 64.3 % compared to the WT (vesicles / 20 μm^2^ / min; GluA1-WT: 13.99 +/− 1.52, AA: 4.99 +/− 1.38) (Fig. 6B, C, Sup. Fig. 6A). The mean speed is significantly reduced for the AA mutant compare to the WT (μm / sec: WT: 1.37 +/− 0.06, AA: 0.92 +/− 0.05) (Fig. 6C, Sup. Fig. 6B). This difference in speed has for consequence that the time spent in the moving state is decreased for the AA mutant to the benefit of the time in pause (Move: WT: 71.68 +/− 0.97 %, AA: 67.22 +/− 1.57 %; pause: WT: 28.08 +/− 0.97 %, AA: 32.78 +/− 1.57 %) (Fig 6C). We analyzed the frequency distribution of speed and found that the pool of receptors going fast (2-4 μm / sec.) is decreased for the AA mutant to the benefit of the slow vesicles (Fig. 6D). This is the case for both OUT and IN directions (Sup. Fig. 6 C). In cLTP condition the binding of 4.1N on the C-ter. domain of GluA1 is necessary for the transport of high velocity vesicles.

Finally, we analyzed the exocytosis of the neosynthetized GluA1 before and after induction of cLTP (Fig. 6 E, F). cLTP increases the exocytosis of GluA1-WT (% of basal; 45 min: 141.06 +/− 7.17; 60 min: 136.52 +/− 7.47) but does not change the localization at the PM of GluA1-AA (% of basal WT; 45 min: before cLTP: 54.84 +/− 3.27, after cLTP: 60.31 +/− 3.70; 60 min before cLTP: 47.18 +/− 4.64, after cLTP: 50.17 +/− 4.04). For this mutant the externalization of GluA1 is decreased both at basal state and inducing cLTP does not change the rate of externalization of the newly synthetized GluA1.

## Discussion

The characteristics of GluA1 IT are very reproducible in basal conditions and heavily regulated during cLTP, presumably allowing for the tuning of newly synthetized AMPAR number at the PM (Hangen et al., 2018). In this study, we analyzed the impact of the expression and binding of 4.1N and SAP97, specifically associated with the intracellular C-ter. domain of GluA1, on the regulation of AMPAR IT (Fig. 7). We detected a differential role between 4.1N or SAP97 binding on GluA1 subunit of AMPAR. These interactions regulate different parameters of IT and exocytosis of neosynthetized GluA1 during basal transmission or during cLTP.

**Figure 7:**
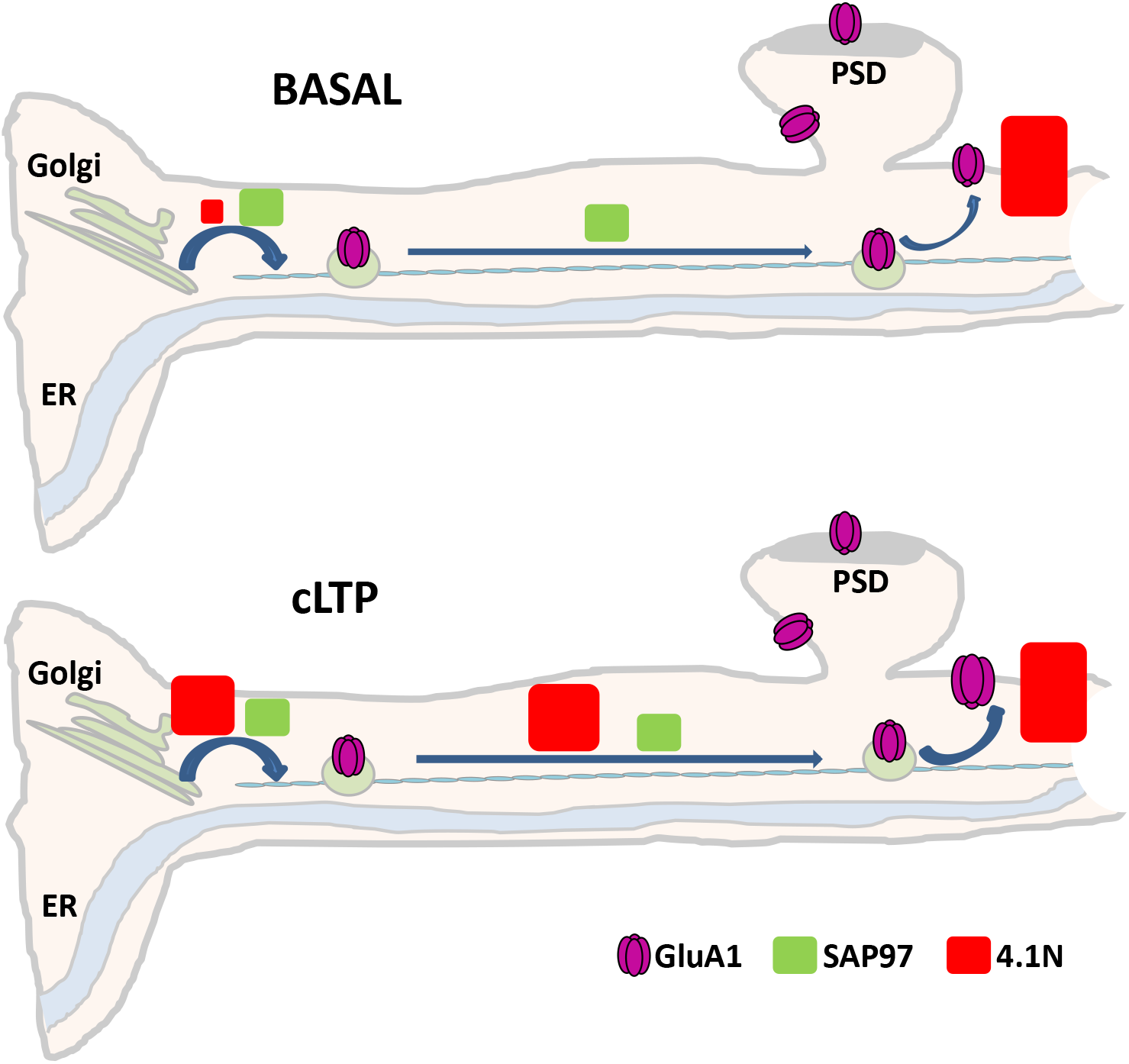
Diagram showing the differential roles of 4.1N/GluA1 and SAP97/GluA1 in basal transmission and during cLTP. In basal condition, upper panel, SAP97 participates to the different phases of IT and, by consequence the number of GluA1 at the PM is decreased. The interaction between 4.1N and GluA1 is fundamental only for the exocytosis of the receptor at the plasma membrane. After induction of cLTP, down panel, SAP97 has the same effect than in basal state. However, 4.1N regulates the exit from the ER/Golgi, the intracellular transport and the exocytosis of GluA1.

Independently decreasing expression of 4.1N or SAP97 inhibited GluA1 IT as well as the exocytosis of the receptor both in basal transmission and after cLTP. For IT regulation, the impact of the reduction of 4.1N was all less drastic than for SAP97. This was the case for the decrease in the number of vesicles released upon addition of AL as well as for the vesicles speed and for the increase in pausing time. In the contrary, we found that the externalization of GluA1 at the PM was equally inhibited by the absence of 4.1N or SAP97.

Whether the C-ter. domain of GluA1 plays any role in synaptic regulation is still uncertain. C-ter. domains of GluA1 and GluA2 are necessary and sufficient to drive NMDA receptor dependent LTP and LTD, respectively (Zhou et al., 2018). Using a single-cell molecular replacement approach and long expression time, the authors found no requirement for the GluA1 C-ter. domain of GluA1 for basal synaptic transmission or for LTP (Granger, Shi et al., 2013). Other work observed a substantially reduced insertion frequency after expression of SEP-GluA1 and TIRF microscopy when the C-ter. domain was deleted (Lin et al., 2009). In our model, we found that deleting the C-ter. domain of GluA1 inhibits almost completely IT and PM localization of newly synthetized GluA1. However, we could still detect rare vesicles transporting the subunit. Our experiments were performed until 120 minutes after addition of the ligand. We cannot rule out that after 24-48 hours some AMPAR could reach the synapse.

To analyze the impact of the interaction between GluA1 and 4.1N, we used two mutants of GluA1. The first one is a complete deletion of the 4.1N binding site, corresponding to a deletion of 14 amino acids close to the last transmembrane domain of GluA1. With this mutant, during basal transmission, we found a decrease in the number of vesicles transporting GluA1, a normal IT and a decrease of exocytosis of GluA1. These effects could be due to the different length of the C-ter. domain of GluA1 or to the unbinding of 4.1N with GluA1. In order to keep the same size of the C-ter. domain, we performed IT and exocytosis experiments with the GluA1-S816A-S818A (AA) mutant that does not bind 4.1N. With this mutant, during basal transmission, only the exocytosis of the protein is impacted. This shows that the interaction between GluA1 and 4.1N is only necessary for receptor exocytosis, without affecting its IT during basal transmission. The interaction with SAP97 has already a role during the transport of GluA1. After cLTP, we observed a decrease of IT properties when the interaction of 4.1N/GluA1 was abolished. The effect of the SAP97/GluA1 interaction during cLTP stays the same than in basal condition.

Previous studies have provided conflicting data regarding SAP97’s influence on synaptic function. Mutant mice expressing GluA1-Δ7 were found to have normal glutamatergic neurotransmission in hippocampal CA1 pyramidal neurons (Zhou et al., 2008). In another study, SAP97 isoforms appeared to regulate the ability of synapses to undergo plasticity by controlling the surface distribution of AMPA and NMDA receptors (Li, Specht et al., 2011). SAP97 directs GluA1 forward trafficking from the Golgi network to the PM. Myosin VI and SAP97 are thought to form a trimeric complex with GluA1, with SAP97 acting as an adaptor between GluA1 and myosin VI to transport AMPA receptors to the postsynaptic plasma membrane (Wu et al., 2002). SAP97 only interacts with GluA1 early in the secretory pathway during its forward trafficking to the PM, suggesting that SAP97 acts on GluA1 solely before its synaptic insertion and that it does not play a major role in anchoring AMPARs at synapses (Sans et al., 2001). SAP97 has also been shown to be associated with motor proteins such as KIF1B (Mok, Shin et al., 2002). Here we found that SAP97 has a major role in IT of AMPAR. Inhibiting expression of SAP97 decreases IT and externalization of neosynthetized GluA1. This effect remains after induction of cLTP. This result is in accordance with the fact that SAP97 has been already shown to be associated with AMPAR during its biogenesis (Sans et al., 2001).

4.1N may function to confer stability and plasticity to the neuronal membrane via interactions with multiple binding partners, including the spectrin-actin-based cytoskeleton, integral membrane channels and receptors. Phosphorylations at both S_816_ and S_818_ residues regulate activity dependent GluA1 insertion, by affecting the interaction between 4.1N and GluA1. In hippocampal neurons, when 4.1N does not bind GluA1 (expression of GluA1-S_816_A-S_818_A), a substantially lower insertion frequency is detected by TIRF microscopy. During LTP these serines are phosphorylated, thus increasing the binding of 4.1N to GluA1 and exocytosis of GluA1. 4.1N is an important player for the expression of LTP, but doesn’t affect basal synaptic transmission (Lin et al., 2009). With our experiments, we show that in basal condition only the exocytosis of neosynthetized GluA1 is inhibited when it does not bind to 4.1N without affecting its IT. However, during cLTP exocytosis and IT properties of GluA1 are dependent of the binding of 4.1N on GluA1.

We show differential roles of 4.1N/GluA1 and SAP97/GluA1 in basal transmission and during cLTP. In basal condition, upper panel Fig. 7, binding of SAP97 with GluA1 participates to the exit from the ER/Golgi, intracellular transport of GluA1 and, by consequence, the number of GluA1 at the PM is decreased (only by 30% compare to the control condition). The interaction between 4.1N and GluA1 is necessary only for the exocytosis of the receptor at the plasma membrane (60% decrease compare to the control) having no effect on IT. After induction of cLTP, down panel Fig. 7, SAP97 has the same effect than in basal state. However, after cLTP the role of the interaction between 4.1N and GluA1 becomes crucial for all phases from the ER to the externalization of AMPAR. By analyzing the GluA1-S816A-S818A mutant we showed that 4.1N/GluA1 interaction regulates the exit of the receptor from the ER/Golgi (number of vesicles of the GluA1-AA decreased by 67% compare to GluA1-WT), the intracellular transport (speed of the GluA1-AA decreased by 35% compare to GluA1-WT), and the exocytosis (localization at the PM of the GluA1-AA decreased by 63% compare to GluA1-WT).

In this study we uncover a new mechanism of regulation of AMPAR trafficking during basal synaptic transmission and synaptic plasticity. It will be important to decipher if this mechanism still all thru *in vivo* and whether its abnormal regulation is involved in some synaptic disruption.

## Acknowledgments

The microscopy was done in the Bordeaux Imaging Center (BIC), a service unit of the CNRS-INSERM and Bordeaux University, member of the national infrastructure “France BioImaging”. We specifically thank the help of Magalie Mondin and Fabrice Cordelières. We like to thank the IINS Cell Biology facility, especially Emeline Verdier, Christelle Breillat, Sophie Daburon and Nicolas Chevrier, for cell culture and plasmid production. This work is currently supported by funding from the Ministère de l’Enseignement Supérieur et de la Recherche, Centre National de la Recherche Scientifique, ERC Grant #787340 Dyn-Syn-Mem, and the Conseil Régional de Nouvelle Aquitaine.

## Author contributions

C. B. designed, performed videomicroscopy experiments and wrote the paper. J. C. performed molecular biology and biochemistry experiments. N. R. performed molecular biology experiments. D. C. wrote the paper. F.C. contributed to all aspects of the project and wrote the paper. The study was conceived and scientifically directed by FC and DC. Authors read and helped revise the manuscript.

## Competing interests

The authors declare no competing financial or other interests that might be perceived to influence the results and/or discussion reported in this paper.

## Data and materials availability

Data and other resources will be available upon request.

## Figure Legends

**Supplemental Figure1:**
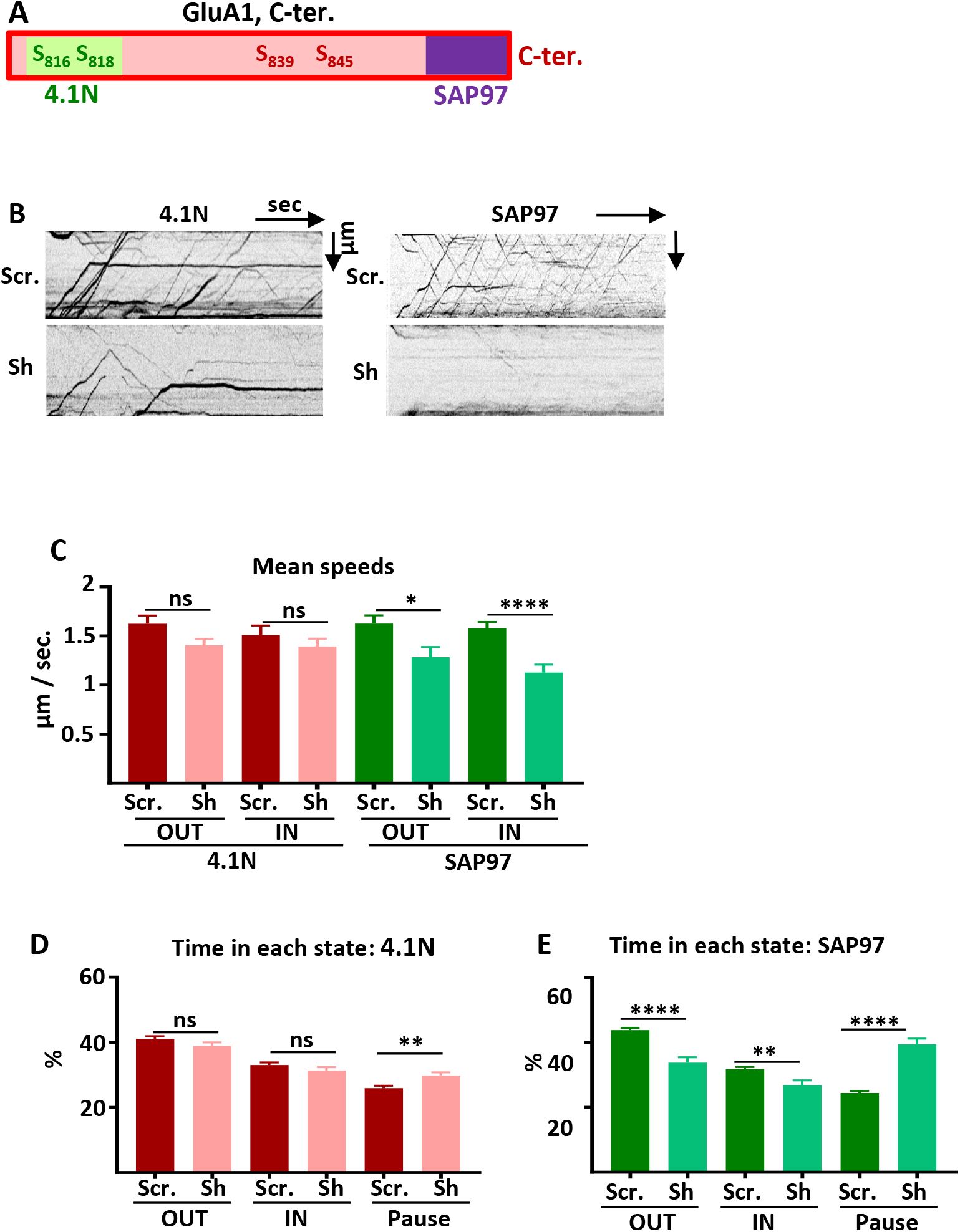
Intracellular transport and exocytosis of GluA1 are regulated by 4.1N and SAP97. **A-** Diagram of the C-terminal (C-ter.) domain of GluA1. **B-** Raw kymographs corresponding to Figure 1D. **C-** Mean speeds OUT and IN of TdTom-GluA1 when expressed with scramble RNA or the corresponding Sh for 4.1N and SAP97. **D-E**-Time spent by a vesicle in each state corresponding to 4.1N (**D**) or to SAP97 (**E**).

**Supplemental Figure2:**
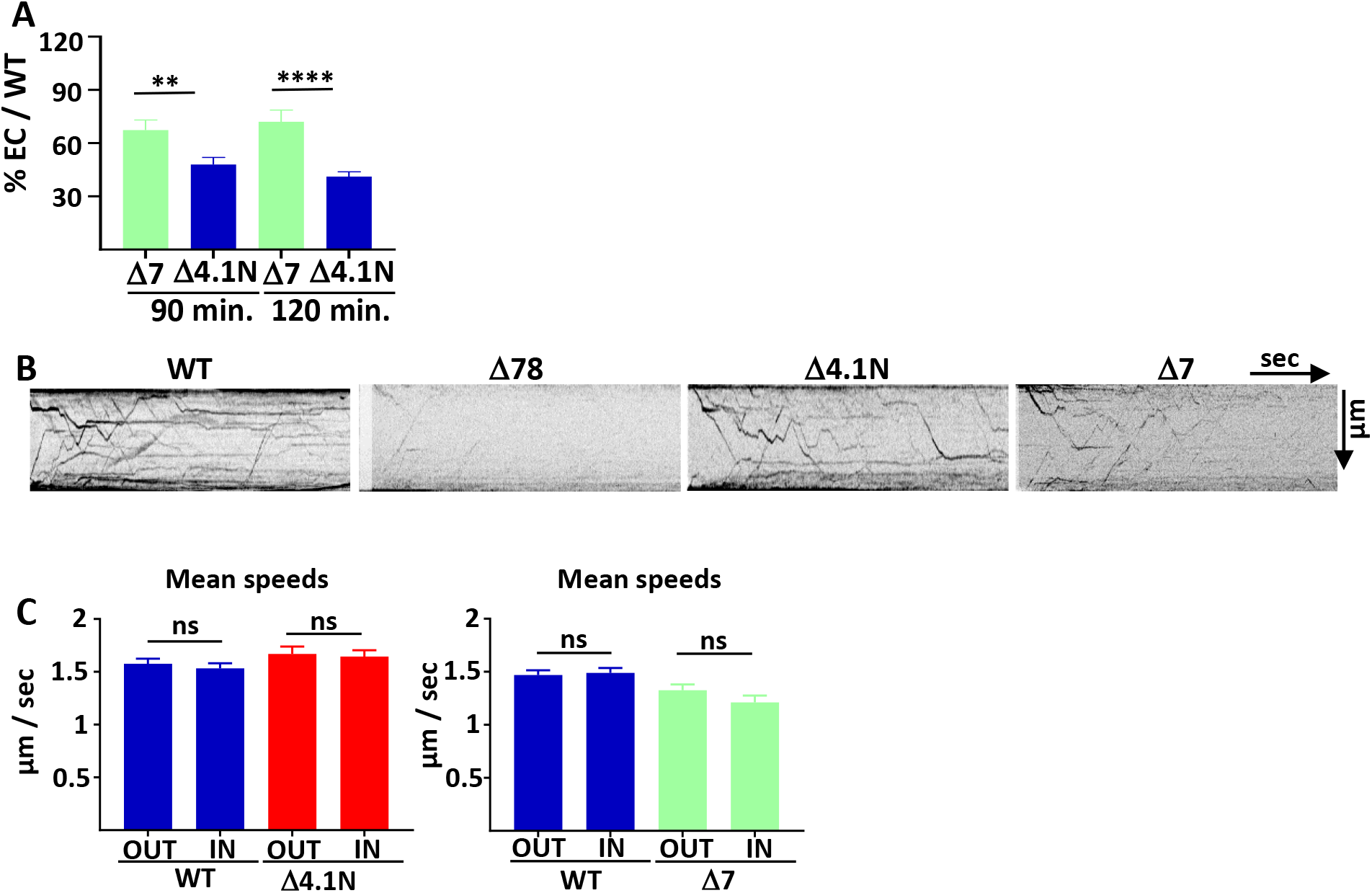
4.1N/GluA1 and SAP97/GluA1 interactions differently regulate GluA1 traffic in basal transmission. **A-** Quantification from Fig. 2E of the surface expression of Δ7 and Δ4.1N at 90 and 120 min. of incubation with AL. The 100 % value correspond to the surface expression of GluA1 WT in the same experiment. **B-** Raw kymographs corresponding to Figure 2F. **C-** Mean speeds OUT and IN of ARIAD-TdTom-GluA1-Δ4.1N (Δ4.1) and ARIAD-TdTom-GluA1-Δ7 (Δ7).

**Supplemental Figure3:**
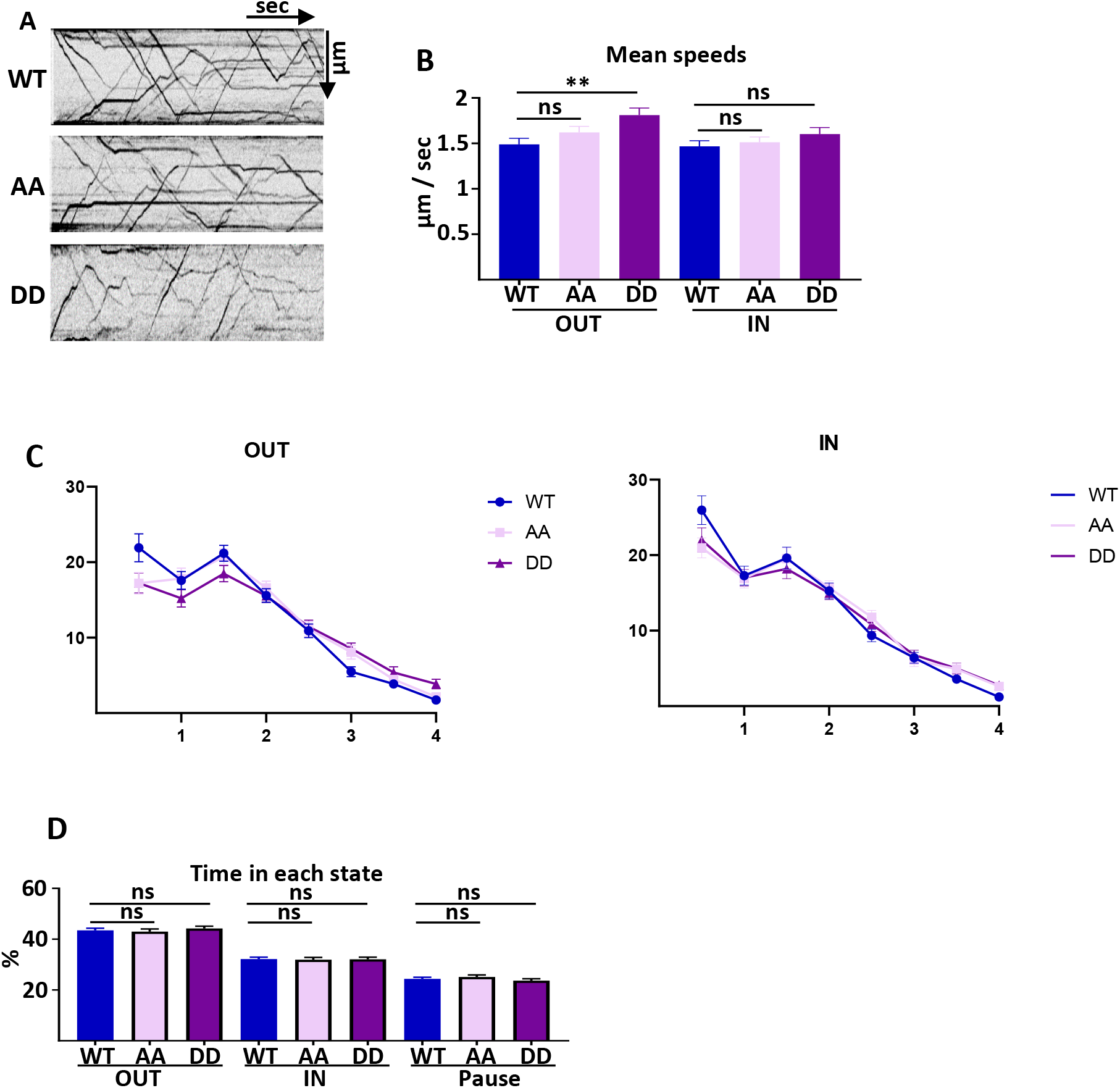
4.1N/GluA1 interaction is only necessary for the exocytosis of GluA1 in basal condition. **A-** Raw kymographs corresponding to Figure 3B. **B-** Mean speeds OUT and IN of ARIAD-TdTom-GluA1-WT (WT), ARIAD-TdTom-GluA1-AA (AA) and ARIAD-TdTom-GluA1-DD (DD). **C-** Frequency distribution of speeds OUT and IN directions for the different mutants as indicated. **D-** Time spent by a vesicle in each state, OUT, IN and Pause.

**Supplemental Fig. 4:**
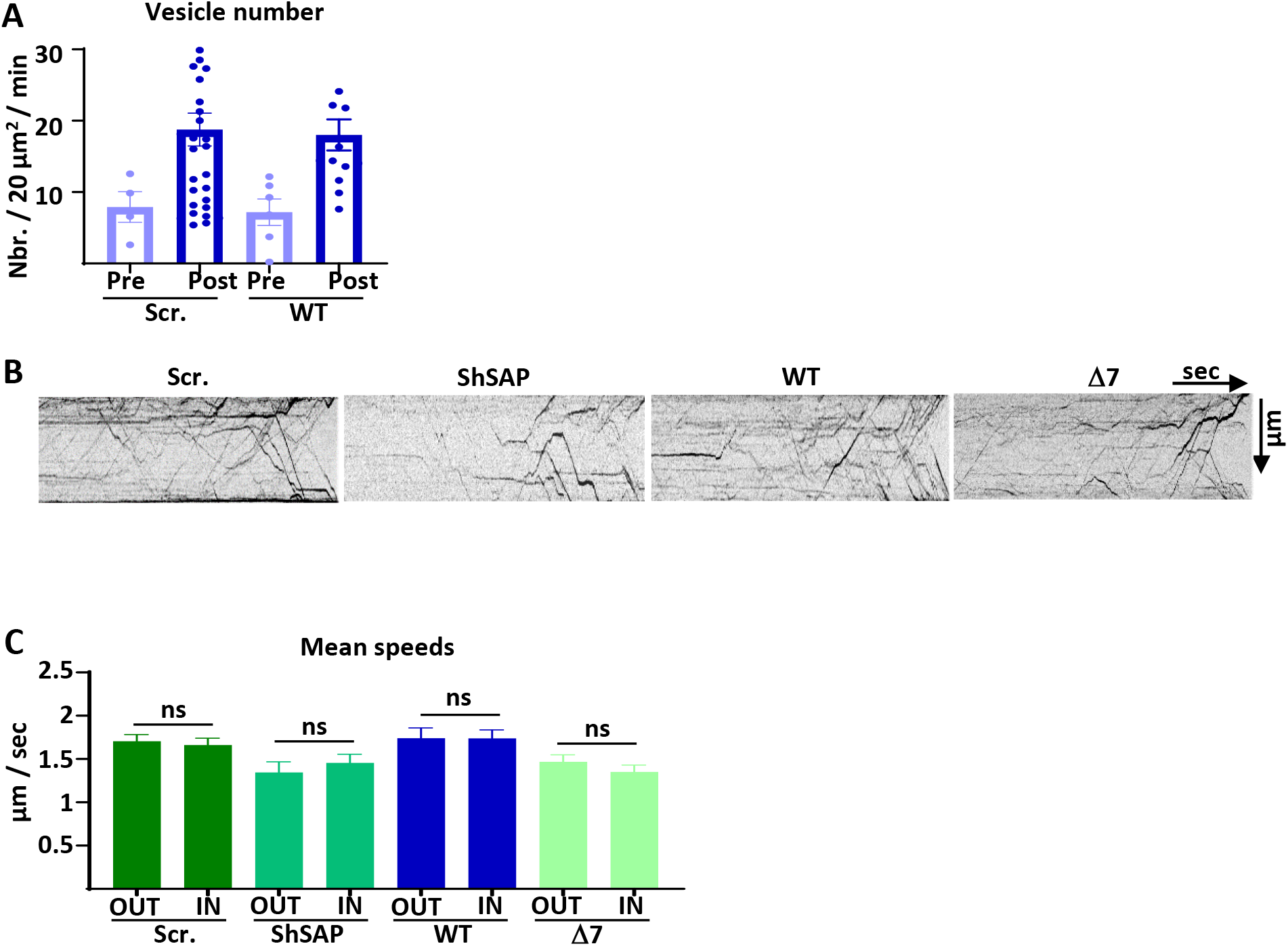
SAP97 and SAP97/GluA1 interaction regulate GluA1 trafficking during cLTP. **A-** Vesicle number before (Pre) and after (Post) cLTP induction for Ariad-TdTom-GluA1-WT with the Scr. RNA against SAP97 or when the GluA1-WT is expressed alone. **B-** Raw kymographs corresponding to Figure 4A. **C-** Mean speeds OUT and IN of the different conditions.

**Supplemental Fig. 5:**
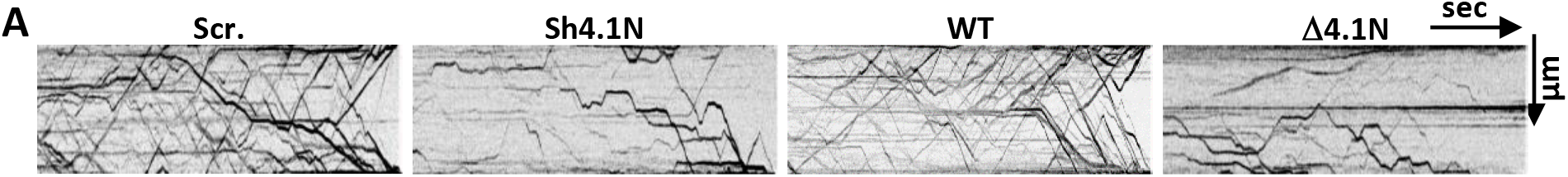
4.1N and 4.1N/GluA1 interaction regulate GluA1 trafficking during cLTP. **A-** Raw kymographs corresponding to Fig. 5A.

**Supplemental Fig. 6:**
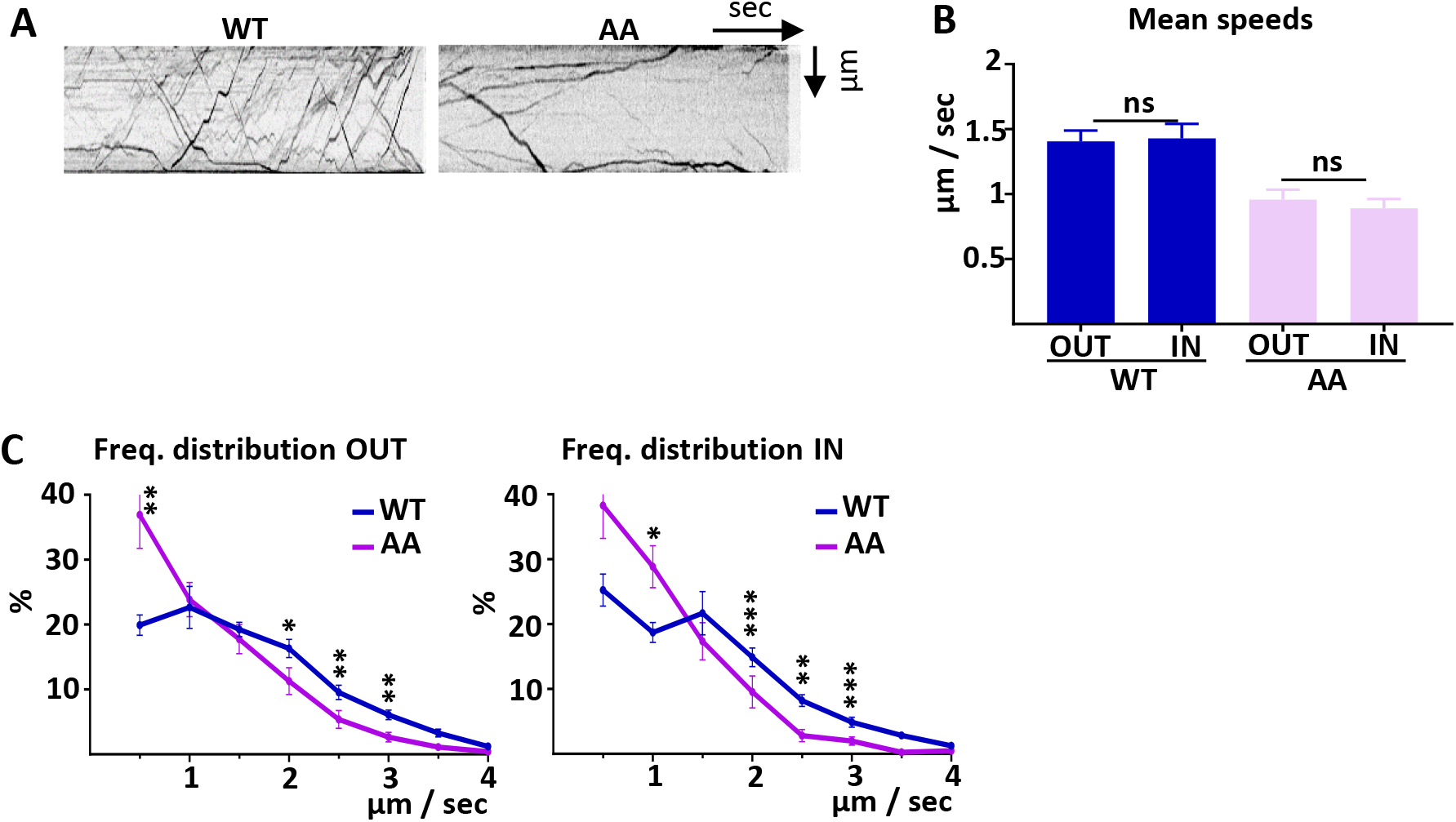
4.1N/GluA1 interaction drives intracellular transport of GluA1 during cLTP. **A-** Raw kymographs corresponding to Fig. 6B. **B-** Mean speeds OUT and IN of the different conditions. **C-** Frequency distribution of speeds OUT and IN for the two proteins WT and AA after induction of cLTP

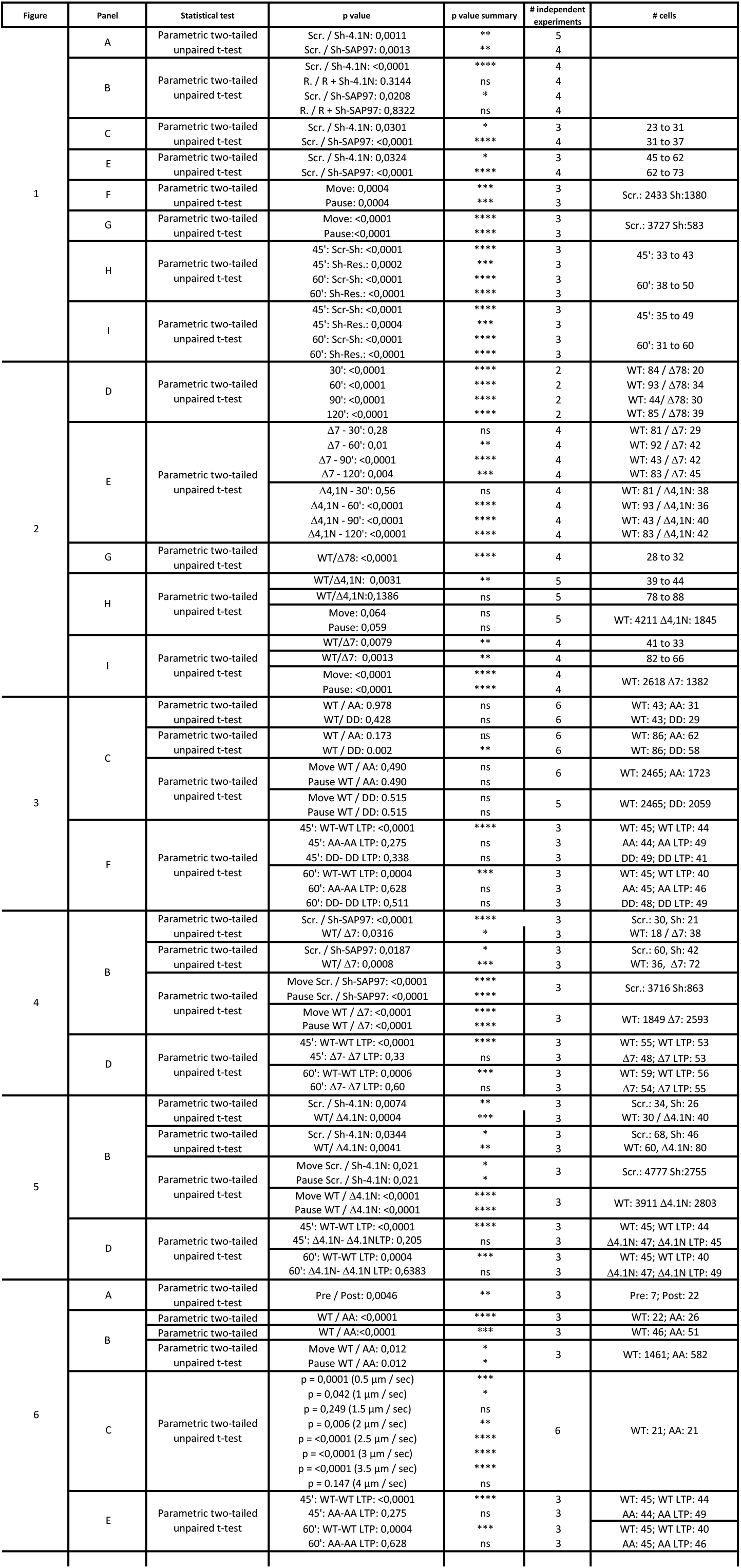

| Figure | Panel | Statistical test | p value | p value summary | # independent experiments | # cells |
| --- | --- | --- | --- | --- | --- | --- |
| 1 | A | Parametric two-tailed unpaired t-test | Scr. / Sh-4.1N: 0,0011<br>Scr. / Sh-SAP97: 0,0013 | **<br>** | 5<br>4 |  |
|  | B | Parametric two-tailed unpaired t-test | Scr. / Sh-4.1N: <0,0001<br>R. / R + Sh-4.1N: 0.3144<br>Scr. / Sh-SAP97: 0,0208<br>R. / R + Sh-SAP97: 0,8322 | ****<br>ns<br>*<br>ns | 4<br>4<br>4<br>4 |  |
|  | C | Parametric two-tailed unpaired t-test | Scr. / Sh-4.1N: 0,0301<br>Scr. / Sh-SAP97: <0,0001 | *<br>**** | 3<br>4 | 23 to 31<br>31 to 37 |
|  | E | Parametric two-tailed unpaired t-test | Scr. / Sh-4.1N: 0,0324<br>Scr. / Sh-SAP97: <0,0001 | *<br>**** | 3<br>4 | 45 to 62<br>62 to 73 |
|  | F | Parametric two-tailed unpaired t-test | Move: 0,0004<br>Pause: 0,0004 | ***<br>*** | 3<br>3 | Scr.: 2433 Sh:1380 |
|  | G | Parametric two-tailed unpaired t-test | Move: <0,0001<br>Pause: <0,0001 | ****<br>**** | 3<br>3 | Scr.: 3727 Sh:583 |
|  | H | Parametric two-tailed unpaired t-test | 45': Scr-Sh: <0,0001<br>45': Sh-Res.: 0,0002<br>60': Scr-Sh: <0,0001<br>60': Sh-Res.: <0,0001 | ****<br>***<br>****<br>**** | 3<br>3<br>3<br>3 | 45': 33 to 43<br>60': 38 to 50 |
|  | I | Parametric two-tailed unpaired t-test | 45': Scr-Sh: <0,0001<br>45': Sh-Res.: 0,0004<br>60': Scr-Sh: <0,0001<br>60': Sh-Res.: <0,0001 | ****<br>***<br>****<br>**** | 3<br>3<br>3<br>3 | 45': 35 to 49<br>60': 31 to 60 |
| 2 | D | Parametric two-tailed unpaired t-test | 30': <0,0001<br>60': <0,0001<br>90': <0,0001<br>120': <0,0001 | ****<br>****<br>****<br>**** | 2<br>2<br>2<br>2 | WT: 84 / Δ78: 20<br>WT: 93 / Δ78: 34<br>WT: 44 / Δ78: 30<br>WT: 85 / Δ78: 39 |
|  | E | Parametric two-tailed unpaired t-test | Δ7 - 30': 0,28<br>Δ7 - 60': 0,01<br>Δ7 - 90': <0,0001<br>Δ7 - 120': 0,004 | ns<br>**<br>****<br>*** | 4<br>4<br>4<br>4 | WT: 81 / Δ7: 29<br>WT: 92 / Δ7: 42<br>WT: 43 / Δ7: 42<br>WT: 83 / Δ7: 45 |
|  |  |  | Δ4,1N - 30': 0,56<br>Δ4,1N - 60': <0,0001<br>Δ4,1N - 90': <0,0001<br>Δ4,1N - 120': <0,0001 | ns<br>****<br>****<br>**** | 4<br>4<br>4<br>4 | WT: 81 / Δ4,1N: 38<br>WT: 93 / Δ4,1N: 36<br>WT: 43 / Δ4,1N: 40<br>WT: 83 / Δ4,1N: 42 |
|  | G | Parametric two-tailed unpaired t-test | WT/Δ78: <0,0001 | **** | 4 | 28 to 32 |
|  | H | Parametric two-tailed unpaired t-test | WT/Δ4,1N: 0,0031<br>WT/Δ4,1N: 0,1386<br>Move: 0,064<br>Pause: 0,059 | **<br>ns<br>ns<br>ns | 5<br>5<br>5<br>5 | 39 to 44<br>78 to 88<br>WT: 4211 Δ4,1N: 1845 |
|  | I | Parametric two-tailed unpaired t-test | WT/Δ7: 0,0079<br>WT/Δ7: 0,0013<br>Move: <0,0001<br>Pause: <0,0001 | **<br>**<br>****<br>**** | 4<br>4<br>4<br>4 | 41 to 33<br>82 to 66<br>WT: 2618 Δ7: 1382 |
| 3 | C | Parametric two-tailed unpaired t-test | WT / AA: 0.978<br>WT / DD: 0.428 | ns<br>ns | 6<br>6 | WT: 43; AA: 31<br>WT: 43; DD: 29 |
|  |  | Parametric two-tailed unpaired t-test | WT / AA: 0.173<br>WT / DD: 0.002 | ns<br>** | 6<br>6 | WT: 86; AA: 62<br>WT: 86; DD: 58 |
|  |  | Parametric two-tailed unpaired t-test | Move WT / AA: 0.490<br>Pause WT / AA: 0.490 | ns<br>ns | 6<br>6 | WT: 2465; AA: 1723 |
|  |  |  | Move WT / DD: 0.515<br>Pause WT / DD: 0.515 | ns<br>ns | 5<br>5 | WT: 2465; DD: 2059 |
|  | F | Parametric two-tailed unpaired t-test | 45': WT-WT LTP: <0,0001<br>45': AA-AA LTP: 0,275<br>45': DD-DD LTP: 0,338<br>60': WT-WT LTP: 0,0004<br>60': AA-AA LTP: 0,628<br>60': DD-DD LTP: 0,511 | ****<br>ns<br>ns<br>***<br>ns<br>ns | 3<br>3<br>3<br>3<br>3<br>3 | WT: 45; WT LTP: 44<br>AA: 44; AA LTP: 49<br>DD: 49; DD LTP: 41<br>WT: 45; WT LTP: 40<br>AA: 45; AA LTP: 46<br>DD: 48; DD LTP: 49 |
| 4 | B | Parametric two-tailed unpaired t-test | Scr. / Sh-SAP97: <0,0001<br>WT/ Δ7: 0,0316 | ****<br>* | 3<br>3 | Scr.: 30, Sh: 21<br>WT: 18 / Δ7: 38 |
|  |  | Parametric two-tailed unpaired t-test | Scr. / Sh-SAP97: 0,0187<br>WT/ Δ7: 0,0008 | *<br>*** | 3<br>3 | Scr.: 60, Sh: 42<br>WT: 36, Δ7: 72 |
|  |  | Parametric two-tailed unpaired t-test | Move Scr. / Sh-SAP97: <0,0001<br>Pause Scr. / Sh-SAP97: <0,0001 | ****<br>**** | 3<br>3 | Scr.: 3716 Sh:863 |
|  |  |  | Move WT / Δ7: <0,0001<br>Pause WT / Δ7: <0,0001 | ****<br>**** | 3<br>3 | WT: 1849 Δ7: 2593 |
|  | D | Parametric two-tailed unpaired t-test | 45': WT-WT LTP: <0,0001<br>45': Δ7- Δ7 LTP: 0,33<br>60': WT-WT LTP: 0,0006<br>60': Δ7- Δ7 LTP: 0,60 | ****<br>ns<br>***<br>ns | 3<br>3<br>3<br>3 | WT: 55; WT LTP: 53<br>Δ7: 48; Δ7 LTP: 53<br>WT: 59; WT LTP: 56<br>Δ7: 54; Δ7 LTP: 55 |
| 5 | B | Parametric two-tailed unpaired t-test | Scr. / Sh-4.1N: 0,0074<br>WT / Δ4.1N: 0,0004 | **<br>*** | 3<br>3 | Scr.: 34, Sh: 26<br>WT: 30 / Δ4.1N: 40 |
|  |  | Parametric two-tailed unpaired t-test | Scr. / Sh-4.1N: 0,0344<br>WT / Δ4.1N: 0,0041 | *<br>** | 3<br>3 | Scr.: 68, Sh: 46<br>WT: 60, Δ4.1N: 80 |
|  |  | Parametric two-tailed unpaired t-test | Move Scr. / Sh-4.1N: 0,021<br>Pause Scr. / Sh-4.1N: 0,021 | *<br>* | 3<br>3 | Scr.: 4777 Sh:2755 |
|  |  |  | Move WT / Δ4.1N: <0,0001<br>Pause WT / Δ4.1N: <0,0001 | ****<br>**** | 3<br>3 | WT: 3911 Δ4.1N: 2803 |
|  | D | Parametric two-tailed unpaired t-test | 45': WT-WT LTP: <0,0001<br>45': Δ4.1N- Δ4.1N LTP: 0,205<br>60': WT-WT LTP: 0,0004<br>60': Δ4.1N- Δ4.1N LTP: 0,6383 | ****<br>ns<br>***<br>ns | 3<br>3<br>3<br>3 | WT: 45; WT LTP: 44<br>Δ4.1N: 47; Δ4.1N LTP: 45<br>WT: 45; WT LTP: 40<br>Δ4.1N: 47; Δ4.1N LTP: 49 |
| 6 | A | Parametric two-tailed unpaired t-test | Pre / Post: 0,0046 | ** | 3 | Pre: 7; Post: 22 |
|  | B | Parametric two-tailed unpaired t-test | WT / AA: <0,0001 | **** | 3 | WT: 22; AA: 26 |
|  |  | Parametric two-tailed unpaired t-test | WT / AA: <0,0001 | *** | 3 | WT: 46; AA: 51 |
|  |  | Parametric two-tailed unpaired t-test | Move WT / AA: 0,012<br>Pause WT / AA: 0,012 | *<br>* | 3<br>3 | WT: 1461; AA: 582 |
|  | C | Parametric two-tailed unpaired t-test | p = 0,0001 (0.5 μm / sec)<br>p = 0,042 (1 μm / sec)<br>p = 0,249 (1.5 μm / sec)<br>p = 0,006 (2 μm / sec)<br>p = <0,0001 (2.5 μm / sec)<br>p = <0,0001 (3 μm / sec)<br>p = <0,0001 (3.5 μm / sec)<br>p = 0.147 (4 μm / sec) | ***<br>*<br>ns<br>**<br>****<br>****<br>****<br>ns | 6 | WT: 21; AA: 21 |
|  | E | Parametric two-tailed unpaired t-test | 45': WT-WT LTP: <0,0001<br>45': AA-AA LTP: 0,275<br>60': WT-WT LTP: 0,0004<br>60': AA-AA LTP: 0,628 | ****<br>ns<br>***<br>ns | 3<br>3<br>3<br>3 | WT: 45; WT LTP: 44<br>AA: 44; AA LTP: 49<br>WT: 45; WT LTP: 40<br>AA: 45; AA LTP: 46 |

## Notes

### Competing Interest Statement

The authors have declared no competing interest.

## References

Bissen D, Foss F, Acker-Palmer A (2019) AMPA receptors and their minions: auxiliary proteins in AMPA receptor trafficking. Cell Mol Life Sci 76: 2133–2169

Cai C, Coleman SK, Niemi K, Keinanen K (2002) Selective binding of synapse-associated protein 97 to GluR-A alpha-amino-5-hydroxy-3-methyl-4-isoxazole propionate receptor subunit is determined by a novel sequence motif. J Biol Chem 277: 31484–90

Choquet D, Triller A (2013) The dynamic synapse. Neuron 80: 691–703

Diaz-Alonso J, Morishita W, Incontro S, Simms J, Holtzman J, Gill M, Mucke L, Malenka RC, Nicoll RA (2020) Long-term potentiation is independent of the C-tail of the GluA1 AMPA receptor subunit. Elife 9

Diaz-Alonso J, Nicoll RA (2021) AMPA receptor trafficking and LTP: Carboxy-termini, amino-termini and TARPs. Neuropharmacology 197: 108710

Diering GH, Huganir RL (2018) The AMPA Receptor Code of Synaptic Plasticity. Neuron 100: 314–329

Fourie C, Li D, Montgomery JM (2014) The anchoring protein SAP97 influences the trafficking and localisation of multiple membrane channels. Biochim Biophys Acta 1838: 589–94

Granger AJ, Shi Y, Lu W, Cerpas M, Nicoll RA (2013) LTP requires a reserve pool of glutamate receptors independent of subunit type. Nature 493: 495–500

Groc L, Choquet D (2020) Linking glutamate receptor movements and synapse function. Science 368

Hangen E, Cordelieres FP, Petersen JD, Choquet D, Coussen F (2018) Neuronal Activity and Intracellular Calcium Levels Regulate Intracellular Transport of Newly Synthesized AMPAR. Cell Rep 24: 1001–1012 e3

Hiester BG, Becker MI, Bowen AB, Schwartz SL, Kennedy MJ (2018) Mechanisms and Role of Dendritic Membrane Trafficking for Long-Term Potentiation. Front Cell Neurosci 12: 391

Hoerndli FJ, Wang R, Mellem JE, Kallarackal A, Brockie PJ, Thacker C, Madsen DM, Maricq AV (2015) Neuronal Activity and CaMKII Regulate Kinesin-Mediated Transport of Synaptic AMPARs. Neuron

Jeyifous O, Waites CL, Specht CG, Fujisawa S, Schubert M, Lin EI, Marshall J, Aoki C, de Silva T, Montgomery JM, Garner CC, Green WN (2009) SAP97 and CASK mediate sorting of NMDA receptors through a previously unknown secretory pathway. Nat Neurosci 12: 1011–9

Kaech S, Banker G (2006) Culturing hippocampal neurons. Nat Protoc 1: 2406–15

Kay Y, Tsan L, Davis EA, Tian C, Decarie-Spain L, Sadybekov A, Pushkin AN, Katritch V, Kanoski SE, Herring BE (2022) Schizophrenia-associated SAP97 mutations increase glutamatergic synapse strength in the dentate gyrus and impair contextual episodic memory in rats. Nat Commun 13: 798

Klopfenstein DR, Vale RD, Rogers SL (2000) Motor protein receptors: moonlighting on other jobs. Cell 103: 537–40

Leonard AS, Davare MA, Horne MC, Garner CC, Hell JW (1998) SAP97 is associated with the alpha-amino-3-hydroxy-5-methylisoxazole-4-propionic acid receptor GluR1 subunit. J Biol Chem 273: 19518–24

Li D, Specht CG, Waites CL, Butler-Munro C, Leal-Ortiz S, Foote JW, Genoux D, Garner CC, Montgomery JM (2011) SAP97 directs NMDA receptor spine targeting and synaptic plasticity. J Physiol 589: 4491–510

Lin DT, Makino Y, Sharma K, Hayashi T, Neve R, Takamiya K, Huganir RL (2009) Regulation of AMPA receptor extrasynaptic insertion by 4.1N, phosphorylation and palmitoylation. Nat Neurosci 12: 879–87

Mandal M, Wei J, Zhong P, Cheng J, Duffney LJ, Liu W, Yuen EY, Twelvetrees AE, Li S, Li XJ, Kittler JT, Yan Z (2011) Impaired alpha-amino-3-hydroxy-5-methyl-4-isoxazolepropionic acid (AMPA) receptor trafficking and function by mutant huntingtin. J Biol Chem 286: 33719–28

Mok H, Shin H, Kim S, Lee JR, Yoon J, Kim E (2002) Association of the kinesin superfamily motor protein KIF1Balpha with postsynaptic density-95 (PSD-95), synapse-associated protein-97, and synaptic scaffolding molecule PSD-95/discs large/zona occludens-1 proteins. J Neurosci 22: 5253–8

Plant K, Pelkey KA, Bortolotto ZA, Morita D, Terashima A, McBain CJ, Collingridge GL, Isaac JT (2006) Transient incorporation of native GluR2-lacking AMPA receptors during hippocampal long-term potentiation. Nat Neurosci 9: 602–4

Rivera VM, Wang X, Wardwell S, Courage NL, Volchuk A, Keenan T, Holt DA, Gilman M, Orci L, Cerasoli F, Jr., Rothman JE, Clackson T (2000) Regulation of protein secretion through controlled aggregation in the endoplasmic reticulum. Science 287: 826–30

Rouach N, Byrd K, Petralia RS, Elias GM, Adesnik H, Tomita S, Karimzadegan S, Kealey C, Bredt DS, Nicoll RA (2005) TARP gamma-8 controls hippocampal AMPA receptor number, distribution and synaptic plasticity. Nat Neurosci 8: 1525–33

Sanderson JL, Gorski JA, Gibson ES, Lam P, Freund RK, Chick WS, Dell’Acqua ML (2012) AKAP150-anchored calcineurin regulates synaptic plasticity by limiting synaptic incorporation of Ca2+-permeable AMPA receptors. J Neurosci 32: 15036–52

Sans N, Racca C, Petralia RS, Wang YX, McCallum J, Wenthold RJ (2001) Synapse-associated protein 97 selectively associates with a subset of AMPA receptors early in their biosynthetic pathway. J Neurosci 21: 7506–16

Schwenk J, Baehrens D, Haupt A, Bildl W, Boudkkazi S, Roeper J, Fakler B, Schulte U (2014) Regional diversity and developmental dynamics of the AMPA-receptor proteome in the mammalian brain. Neuron 84: 41–54

Schwenk J, Boudkkazi S, Kocylowski MK, Brechet A, Zolles G, Bus T, Costa K, Kollewe A, Jordan J, Bank J, Bildl W, Sprengel R, Kulik A, Roeper J, Schulte U, Fakler B (2019) An ER Assembly Line of AMPA-Receptors Controls Excitatory Neurotransmission and Its Plasticity. Neuron 104: 680–692 e9

Shen L, Liang F, Walensky LD, Huganir RL (2000) Regulation of AMPA receptor GluR1 subunit surface expression by a 4. 1N-linked actin cytoskeletal association. J Neurosci 20: 7932–40

Walensky LD, Blackshaw S, Liao D, Watkins CC, Weier HU, Parra M, Huganir RL, Conboy JG, Mohandas N, Snyder SH (1999) A novel neuron-enriched homolog of the erythrocyte membrane cytoskeletal protein 4.1. J Neurosci 19: 6457–67

Wu H, Nash JE, Zamorano P, Garner CC (2002) Interaction of SAP97 with minus-end-directed actin motor myosin VI. Implications for AMPA receptor trafficking. J Biol Chem 277: 30928–34

Zhou W, Zhang L, Guoxiang X, Mojsilovic-Petrovic J, Takamaya K, Sattler R, Huganir R, Kalb R (2008) GluR1 controls dendrite growth through its binding partner, SAP97. J Neurosci 28: 10220–33

Zhou Z, Liu A, Xia S, Leung C, Qi J, Meng Y, Xie W, Park P, Collingridge GL, Jia Z (2018) The C-terminal tails of endogenous GluA1 and GluA2 differentially contribute to hippocampal synaptic plasticity and learning. Nat Neurosci 21: 50–62

